# Biofilms and core pathogens shape the tumour microenvironment and immune phenotype in colorectal cancer

**DOI:** 10.1101/2023.10.20.563034

**Authors:** Lasse Kvich, Blaine Gabriel Fritz, Henrike Zschach, Thilde Terkelsen, Hans Raskov, Kathrine Høst-Rasmussen, Morten Ragn Jakobsen, Alexandra Gabriella Gheorghe, Ismail Gögenur, Thomas Bjarnsholt

**Affiliations:** Center for Surgical Science, Department of Surgery, Zealand University Hospital, Køge, Region Zealand, 4690, Denmark; Costerton Biofilm Center, Department of Immunology and Microbiology, University of Copenhagen, Copenhagen, The Capital Region, 2200, Denmark; Center for Health Data Science, University of Copenhagen, Copenhagen, The Capital Region, 2200, Denmark; Department of Forensic Medicine, Faculty of Health and Medical Sciences, University of Copenhagen, Copenhagen, The Capital Region, 2100, Denmark; V 2200, Copenhagen, Denmark; Department of Clinical Microbiology, Rigshospitalet, Copenhagen, The Capital Region, 2100, Denmark

**Author notes:** Corresponding authors: Thomas Bjarnsholt **&** Ismail Gögenur. These authors contributed equally to this work, and the co-first authors agreed upon the order (Lasse Kvich, Blaine Gabriel Fritz). Senior author (Ismail Gögenur, Thomas Bjarnsholt).

**Keywords:** colorectal cancer (CRC), biofilms, *Fusobacterium nucleatum*, *Bacteroides fragilis*, *In Situ* Hybridization, Fluorescence, Sequence Analysis, RNA

## Abstract

**Objective:** Growing evidence links bacterial dysbiosis with colorectal cancer (CRC) carcinogenesis, characterized by an increased presence of core pathogens such as *Bacteroides fragilis* and *Fusobacterium nucleatum*. Here, we characterized the *in situ* biogeography and transcriptional interactions between bacteria and the host in mucosal colon biopsies.

**Design:** The influence of CRC core pathogens and biofilms on the tumour microenvironment (TME) was investigated in biopsies from patients with and without CRC (paired normal tissue and healthy tissue biopsies) using fluorescence *in situ* hybridization and dual-RNA sequencing.

**Results:** Tissue-invasive, mixed-species biofilms enriched for *B. fragilis* and *F. nucleatum* were observed in CRC tissue, especially in right-sided tumours. *Fusobacterium spp.* was associated with increased bacterial biomass and inflammatory response in CRC samples. CRC samples with high bacterial activity demonstrated increased expression of pro-inflammatory cytokines, defensins, matrix-metalloproteases, and immunomodulatory factors. In contrast, the gene expression profiles of CRC samples with low bacterial activity resembled healthy tissue samples. Moreover, immune cell profiling showed that *B. fragilis* and *F. nucleatum* modulated the TME and correlated with increased infiltration of neutrophils and CD4^+^ T-cells. Overall, bacterial activity was critical for the immune phenotype and correlated with the infiltration of several immune cell subtypes, including M2 macrophages and regulatory T-cells.

**Conclusion:** Biofilms and core pathogens shape the TME and immune phenotype in CRC. Our results support that *Fusobacterium spp*. may provide a future therapeutic target to reduce biofilms and the inflammatory response in the TME while highlighting the importance of widening the scope of bacterial pathogenesis in CRC beyond core pathogens.

## INTRODUCTION

Several studies demonstrate associations between altered gut microbiota composition and colorectal cancer (CRC) ^1–3^. This imbalance in the gut microbiota is termed microbial dysbiosis, and it is considered to contribute to CRC pathogenesis ^4^ ^5^. Microbial dysbiosis allows opportunistic bacteria, such as *Bacteroides fragilis* and *Fusobacterium nucleatum*, to accumulate and increase inflammation, a known risk factor for CRC ^6^ ^7^. The mechanisms whereby bacteria accelerate carcinogenesis have been explored in numerous *in vitro* and animal studies, where specific bacterial species influence pathways related to the initiation and progression of CRC ^8^. Specifically, *B. fragilis* and *F. nucleatum* can alter the tumour microenvironment (TME) and facilitate CRC progression by fueling an inflammatory response ^9^ ^10^ and suppressing anti-tumourigenic immune cells ^11–13^. However, mechanistic insights from animal experiments or *in vitro* studies do not adequately characterize the bacteria and host interplay ^14^. Recently, metatranscriptomic studies have investigated the bacteria and host interplay in infectious diseases ^15^ ^16^, but metatranscriptomic studies in CRC are still scarce^17^ ^18^.

In this study, we characterize the interplay between the host and active mucosa-associated bacteria in CRC, paired normal, and healthy tissue using fluorescence *in situ* hybridization and dual-RNA sequencing. Species-specific and pan-microbial microscopic examination of cross-sectioned whole biopsies revealed the involvement of *Fusobacterium spp.* in the accumulation of bacterial biomass (biofilms) and acute inflammation in CRC samples. Further, immune cell profiling revealed that overall bacterial activity and *B. fragilis* and *F. nucleatum* activity correlated with the infiltration of specific immune cells. Our findings show that biofilms and CRC core pathogens fuel the inflammatory response in the TME and suggest *Fusobacterium spp*. as a therapeutic target to reduce inflammation-driven CRC carcinogenesis.

## MATERIALS AND METHODS

### Ethics and patient recruitment

Patients were recruited at Zealand University Hospital from December 2020 to December 2021. Inclusion criteria were persons >18 years old with written, approved consent admitted for a colonoscopy exam or patients undergoing surgery. Subjects were divided into patients with pathologically verified CRC (any T-stage) and healthy persons with no underlying gastrointestinal malignancies or diseases. No other inclusion or exclusion criteria were used. This study was approved by the Danish Regional Ethical Committee (SJ-826) and the Danish Data Protection Agency (REG-024-2020).

### Study design and sampling

Mucosal colon biopsies were collected from 40 patients with pathologically verified CRC (any T-stage) for RNA sequencing and microscopy. Biopsies were collected from the tumour and paired normal tissue >10 cm away from the tumour, if possible. Mucosal colon biopsies were also collected from 40 individuals with no observed gastrointestinal diseases as healthy control samples. Biopsies were sampled from anatomically different areas of the colon (right and left-sided). An even amount of left and right-sided samples was ensured between patients with CRC and individuals with no observed gastrointestinal diseases. A pathologist screened CRC biopsies to ensure the presence of carcinoma. After collection, biopsies for RNA sequencing were placed immediately in RNAlater® (Invitrogen, MA, USA) and stored for a minimum of 18-24 hours at 5°C before further processing. Samples were centrifuged at 3000 x g for 5 minutes, the RNAlater® was removed, and the tissue was stored at −80°C until RNA purification. For microscopy analysis, two biopsies were collected per person (CRC and healthy) and area of sampling (tumour and paired normal tissue). Biopsies were fixated immediately in 4% buffered paraformaldehyde (pH 7.4) and stored at 4° C for at least 24 hours before being embedded in paraffin (FFPE).

### FISH probes

Peptide Nucleic Acid (PNA) probes targetting *B. fragilis* (Bfrag-998) ^19^ and *F. nucleatum* (FUS714) ^20^ were ordered from Biomers (Ulm, DE). Bfrag-998 was tagged at the 5’end with Cyanine5 (Cy5-5’-GTTTCCACATCATTCCACTG-’3) and FUS714 was tagged at the 5’end with Cyanine3 (Cy3-5’-GGCTTCCCCATCGGCATT-’3). A universal (BacUni) bacterial probe (AdvanDx, Woburn, MA) tagged at the 5’end with Texas Red was used to visualize all bacteria. The specificity of Bfrag-998 and FUS714 was checked with the Basic Local Alignment Search Tool (BLAST) function in the NCBI database. The FUS714 probe aligned with other *Fusobacterium* strains, including four subspecies of *Fusobacterium nucleatum* (polymorphum, nucleatum, vincentii, and animalis), and it was also complementary to two other bacterial species belonging to the Fusobacteriaceae family (*Ilyobacter polytropus* and *Propionigenium modestum*); thus microscopy findings are referred to as *Fusobacterium spp*. The Bfrag-998 probe was specific for *B. fragilis*.

### Peptide nucleic acid fluorescence *in situ* hybridization (PNA-FISH)

FFPE samples were sectioned onto glass slides (3-5 µm sections) before standard xylene-deparaffinization. Each glass slide had two sections with two biopsies per person. PNA-FISH was carried out according to a standard in-house protocol with few adjustments ^21^. The hybridization buffer was prepared according to Stender *et* al. ^22^ with a final concentration of 250 nM for each probe. Samples were covered with 30 µL hybridizations buffer with a mixture of either all three probes or the specific probes (Bfrag-998 and FUS714) and left for incubation for one and a half hours at 56° C. Samples were then washed in a pre-warmed (56° C) washing buffer (AdvanDx, USA) for 30 minutes and left to dry for 15 minutes. Samples were counterstained with 0.3 µM 4′,6-diamidino-2-phenylindole (DAPI; Life Technologies, OR, USA) for 15 minutes before rinsing with cold phosphate-buffered saline pH = 7.5 (Panum Institute Substrate Department, University of Copenhagen, DK). The samples were left to dry before an antifade reagent was applied (ProLong^TM^ Gold, Thermo Fisher Scientific, UK). Finally, a cover glass (Marienfeld, DE) was added and sealed with clear nail polish.

### Microscopy and image processing

All fluorescence microscopy was performed on an inverted Zeiss LSM 880 confocal microscope (Zeiss, Jena, DE), using either a Plan-Apochromat 63x/1.40 Oil DIC M27 objective or an EC Plan-Neofluar 40x/1.30 Oil DIC M27 objective. All images were obtained in 16-bit, and the instrument automatically assigned the optimal pinhole size (Airy units) for each analysis described below. Illustrative images were processed with maximum intensity projection and a Gaussian filter for smoothing, and the brightness was increased by 20% for presentation purposes. All illustrative images were processed in the Imaris 9.7.2 software (Oxford Instruments, UK), either as 3D projections (bacterial biomass) or 2D projections (illustrative images). In addition, pseudo-colouring was used for representative images to separate different bacterial populations from each other. *Fusobacterium spp*. were pseudo-coloured green, *B. fragilis* red, and all other bacteria purple.

### Bacterial biomass

A 594 nm laser was used to excite the BacUni probe for bacterial biomass imaging, and the fluorescence emission was detected in 597–661 nm intervals. One biopsy was selected for analysis in the different groups (primary tumour, paired normal, and healthy tissue). The first biopsy encountered during microscopy was used for this analysis to avoid selection bias. Area (tile scan) and depth (z-stack) were manually set for each biopsy. Biomass (µm3) was measured as previously described ^23^, using the Measurement Pro addon in Imaris (Oxford Instruments, UK). Imaris utilizes a pixel quantitative approach to measure biomass, where a mask was created for total biomass (tissue and bacteria) and bacterial biomass (only bacteria) according to the thresholding of fluorescence intensity.

### Probe validation and *B. fragilis* and *Fusobacterium spp.* prevalence

The FUS714 and Bfrag-998 probes were qualitatively validated to ensure correct differentiation during microscopy. The differentiation of *Fusobacterium nucleatum ssp. nucleatum* (ATCC 25586) and *Bacteroides fragilis* (ATCC 25285) were tested on spiked lung tissue explanted from a mink (surplus material from animal studies). Overnight cultures with both strains were prepared and grown in brain-heart infusion media (Sigma-Aldrich, USA) under anoxic conditions for 24 hours at 37° C, and tissue was subsequently spiked in a 1:1 ratio by injection. Afterwards, the tissue was fixated in 4% buffered paraformaldehyde, paraffin-embedded, and treated according to the PNA-FISH method described above. A 561 and 633 nm laser was used for the excitation of Cy3 (FUS714) and Cy5 (Bfrag-998), respectively. Fluorescence emission was detected in 549–573 nm intervals for FUS714 and 632-705 nm intervals for Bfrag-998. After initial testing and adjustment on spiked tissue, the probes were tested *ex vivo* on tumour biopsies with similar settings to ensure the correct differentiation of bacterial populations. Two sections with two biopsies were screened per person in each group to assess the prevalence of *B. fragilis* and *Fusobacterium spp*. Excitation of DAPI was acquired at 405 nm, and the fluorescence emission was detected in 415–488 nm intervals. The settings described above were used for FUS714 and Bfrag-998. The narrow intervals for FUS714 were used to avoid background fluorescence from the tissue. Sequential multiple-channel fluorescence scanning was used to avoid or reduce bleed-through (cross talk) across the fluorophores. All findings (scattered cells or aggregated bacteria) were counted and included in the prevalence measurement. In addition, the biomass was assessed for *B. fragilis* and *Fusobacterium spp.* on a subset of samples (n = 7) using the same excitation and emission intervals described above.

### Histopathology and inflammation score

Two pathologists evaluated the histopathology and scored inflammation in CRC samples to assess whether bacterial biomass (biofilms), *Fusobacterium spp.*, or *B. fragilis* affected the degree of inflammation. Tissue sections were cut at 4 µm thicknesses, mounted on glass slides, and stained with Hematoxylin & Eosin. The degree of inflammation was scored 0 (no inflammation), 1 (mild inflammation), 2 (moderate inflammation), and 3 (severe inflammation) ^24^ ^25^. A score was given for acute and chronic inflammation, reflecting the infiltration of polymorphonuclear leukocytes (PMNs) and lymphocytes, respectively. Two pathologists performed the histological assessment in a blinded way. Histological analysis was performed using a Leica DM 4000 B LED light microscope. In addition, the co-localization of bacterial biomass and necrotic tissue was assessed by a pathologist in samples with high bacterial biomass (n = 21) using an EVOS M7000 microscope (Thermofisher MA, USA).

### RNA extraction and purification

Biopsies were removed from −80°C and placed immediately into 2 mL microtubes (Sarstedt, Nuembrecht, Germany) filled to ∼1/3 volume with 2 and 0.1 mm diameter zirconia beads (Biospec, OK, USA) on ice. Eight hundred microliters of ice-cold Trizol (Invitrogen, MA, USA) containing 10 uL/mL β-mercaptoethanol (Sigma-Aldrich, MO, USA) was added to each tube. Samples were homogenized 3 x 30s at 7000 power in a MagnaLyzer® (Roche Diagnostics, Basel, Schweiz) and placed on ice for ∼1 minute between each homogenization. One-hundred sixty microliters of chloroform (Sigma-Aldrich, MO, USA) was added, and the tubes were shaken by hand for 45 s. Samples were spun down at 13.000 x g at 4°C for 15 min. The aqueous phase was collected in a 1.5 mL Eppendorf tube. Four hundred microliters of cold isopropanol (Sigma-Aldrich, MO, USA) and 2uL of linear acrylamide (ThermoFisher, MA, USA) were added to each sample. Tubes were then inverted 4-6 times and incubated at −20 °C for 60-90 minutes. Samples were spun down again as previously, and the supernatant was removed. The pellet was washed twice with 900uL of freshly prepared and ice-cold 80 % ethanol. After the second wash, the ethanol was removed, and the samples were air-dried for ∼5-10 minutes to evaporate excess ethanol. The pellet was then resuspended in 20 uL of nuclease-free water. The concentration and purity of extracted RNA were assessed with a Nanodrop spectrophotometer (ThermoFisher, MA, USA). The purified RNA was stored at −80°C.

### Ribosomal RNA depletion

Ribosomal RNA (rRNA) depletion was performed using the riboPOOL™ kit (siTOOls Biotech, Germany). One microgram of purified RNA was used as input, if available. If one microgram in 15uL water was not possible due to low concentration, 15 uL of the purified RNA was used as input. The protocol was performed as described in the riboPOOLKitManual_V1.3. The riboPOOL used for the depletion was a 100:1 combination of the Human riboPOOL (riboPOOL_054) and Pan-Prokaryote riboPOOL (riboPOOL_003). Eighty microliters of rRNA-depleted RNA were treated with RQ1 RNAse-free DNAse (Promega, USA) (10uL DNAse + 10uL buffer) per sample and incubated for 30 min at 37°C. The rRNA-depleted and DNAse-treated RNA was then cleaned with the Zymo RNA Clean and Concentrate-5 kit (Zymo Technologies, USA) and eluted in 8 uL nuclease-free water.

### Library preparation and sequencing

One hundred nanograms of rRNA-depleted, DNAse-treated RNA in 5uL water was used as input to the NEB Ultra II directional library-preparation kit (New England BioLabs, MA, USA). If the concentration was less than this, 5uL of the rRNA-depleted, DNAse-treated RNA was used. The protocol was performed as described in the manual for rRNA-depleted RNA. Ten or twelve PCR cycles were used for the final enrichment step for samples with inputs of 100 ng or less, respectively. Quality and concentration of final libraries were measured by Qubit (1x dsDNA kit; Invitrogen, MA, USA) and Bioanalyzer (DNA High Sensitivity Chip; Agilent, CA, USA). Samples were pooled in equimolar amounts, cleaned with the 1.8x HighPrep™ PCR beads (Magbio, Lusanne, Schweiz), and sequenced on an Illumina NovaSeq 6000 instrument. Samples 1-33 and 34-118 were sequenced in S2, and S4 flow cells, respectively, with v1.5 reagents and 150 PE reads.

### Preliminary processing of raw RNA sequencing data

Raw sequencing data (bcl. files) were demultiplexed into forward and reverse reads for each sample using bcl2fastq v2.20.0 from Illumina and concatenated across lanes. Cutadapt v3.4 ^26^ was used to trim adapters and filter out short reads (maximum error rate = 0.005, minimum length = 33, minimum overlap = 7). rRNA reads were removed with sortmeRNA v4.3.4 ^27^ using all of the included databases. The rRNA-depleted reads were then aligned to the human reference genome (GRCh38.p13, Ensembl release 106, primary assembly, build: GCA_000001405.28) with bwa-mem v0.7.17 ^28^. Reads mapping to annotated, gene-level features were counted with featureCounts (parameters: –p –O –fracOverlap 0.2 -J -t gene) from subread v2.0 ^29^ using the .gtf Ensembl annotations (GRCh38.106, Ensemble release 106). Outputted files were then concatenated by columns into a final gene-count matrix. The trimmed reads were also classified using Kraken v.2.1.2 ^30^ to determine bacterial community composition using the standard RefSeq index database (obtained from: https://benlangmead.github.io/aws-indexes/k2). Abundances of actively transcribing bacteria were estimated with Bracken v2.7 ^31^.

### Bacterial community composition

The bracken output was multiplied by a scaling factor to account for differences in sequencing depth between the two sequencing runs (Samples 1-33 and 34-118). This scaling factor was calculated as the number of reads in the sample with the lowest number divided by the number of reads in a given sample. Scaled counts were used when comparing across the samples, and unscaled counts were used when comparing within-sample variation. Also, a threshold was applied to remove noise from low-abundance taxa. The cut-off was determined by visual inspection of the log-10 transformed scaled counts distribution, and the intersection between the two independent, overlapping, normal-distributed populations was used as the cut-off (Figure S1); thus, all scaled counts < 0.125 were set to 0. Differences in mean scaled counts of bacteria, *F. nucleatum*, and *B. fragilis* between conditions (CRC vs. healthy, CRC vs. paired) were tested with a Wilcoxon rank-sum test or Wilcoxon signed-rank tests for paired and independent samples.

### Differential Gene Expression and Functional Enrichment Analysis

Differential gene expression and functional enrichment analyses were performed to identify differentially expressed genes (DEGs) and pathways between CRC and Healthy/Paired-normal tissue. First, the count data were filtered to include only protein-coding transcripts using the biomaRt package in R ^32^. A dummy variable (“cancer”) for samples originating from CRC (“CRC”) or Healthy/Paired-normal (“no_CRC”) was encoded. Differential gene expression analysis was performed using the DESeq function from DEseq2 v1.36.0 with default settings and the formula “∼ cancer”. DEGs with an adjusted p-value less than 0.05 and |log2 fold-change|>2 were used for further analysis. We adapted a previously published approach to test whether differentially expressed genes represented an enrichment of known biological pathways ^33^. Briefly, the Kyoto Encyclopedia of Genes and Genomes (KEGG), Pathway Interaction Database (PID), and REACTOME (a database of reactions, pathways, and biological processes) canonical data sets were downloaded from the MsigDB database. Pathways with a minimum of 25 and maximum of 85 genes and overlap of at least five genes with the DEGs of interest were included. A Fisher’s exact test was performed for each pathway and the resulting p-values were adjusted with Benjamini-Hochberg correction. This analysis was performed separately for genes showing positive or negative log2 fold changes, respectively. To analyze the effect bacteria on host gene expression in tumour tissue, the count matrix described above was further subsetted to include only CRC samples. A binary variable was then created to identify samples containing a high and low bacterial signal (defined as outliers on a scaled count, Figure S1). Differential gene expression and functional enrichment analysis were then performed as described above with this variable instead of the “cancer” variable.

### Immune cell profiling

There are many different methods for estimating immune cell infiltration from RNA sequencing data ^34^ ^35^. Each of these likely return different scores due to the underlying algorithm employed by the method and its predefined immune cell populations. Therefore, a consensus approach was utilized using the R package immunedeconv ^36^. This package implements multiple methods, including quantiseq, epic, estimate, mcp_counter, xcell, consensus_tme, and timer, for estimating sample immune infiltration based on bulk RNA sequencing reads. Filtering and normalization of the raw-count matrix of gene expression values were performed with limma ^37^. Low-expressed genes were filtered out with the filterByExpr function, the scaling factor was estimated with calcNormFactors, and a matrix of TMM (Trimmed Mean of M-values) counts was generated with the cpm function. As each deconvolution method for estimating a score returns different sets/subsets of immune cell types, generalized cell categories were defined for each cell type, e.g., CD4+ and CD8+, and regulatory T-cells were classified as T-cells. Also, given that some methods (quantiseq, epic) return a proportion of a given cell type to all cells while others (mcp_counter, xcell, consensus_tme, timer, estimate) return scores on varying scales, the analyses were performed separated for proportions and scores. Normalization was performed by positive centering of the scores to adjust for differences in scales between methods utilizing a score, adding the smallest observed score value (greater than zero) in each method as a pseudo count to scores that were 0, after which the values were log2 transformed. Finally, heatmaps were generated to visualize the hierarchical clustering of samples according to the immune cell profile score. The R-packages NBclust, cluster, as well as the results from the hierarchical clustering were used to select the optimal number of clusters (n=4). A nonparametric Kruskal-Wallace test was used to test whether the total bacterial, *F. nucleatum*, or *B. fragilis* activity affected the immune cell clusters. Post-hoc, pairwise comparisons between clusters were then performed with a Dunn test.

### Statistics

Colorectal cancer RNAseq data from The Cancer Genome Atlas https://portal.gdc.cancer.gov was used to estimate the required sample sizes. Based on the bacterial gene transcription, which is expectedly lower than the host, it was estimated that 40 patients should be included in each group. An alpha of 0.5 and a power of 0.8 was used. Bacterial biomass (µm^3^) was log-transformed (Log) to ensure normally distributed data unless otherwise stated. In some cases, CRC data were separated into high and low bacterial biomass using the average of all bacterial biomass measurements as the cut-off (4.2 Log um^3^). The CRC sample with the missing paired sample was excluded from all paired analyses. Graphs and statistics were carried out with either GraphPad Prism 9.3.1 (GraphPad Software, La Jolla, California, USA) or the R software v3.6.0 (R Development Core Team 2004). An adjusted p-value was reported in the case of multiple testing, and a p-value ≤ 0.05 was considered significant.

## RESULTS

### Patient characteristics

Standard osmotic bowel prep was used in all cases before sampling, and only two patients with CRC reported using antibiotics before inclusion and sampling. Patients with CRC had a higher ASA score (American Society of Anesthesiologists - a metric to determine if a patient is healthy enough to tolerate surgery and anaesthesia) than healthy persons (Table 1). This difference was expected, given the aetiology of CRC and the demographic characteristics of the CRC population. More males were diagnosed with CRC, and left-sided tumours were more prevalent than right-sided, reflecting the normal distribution of CRC. Biopsies were mainly collected from the rectum and colon sigmoideum (Figure 1A). Due to advanced disease, ten patients did not receive pathologically verified tumour staging; palliative care was provided in these cases rather than surgery.

**Figure 1.**
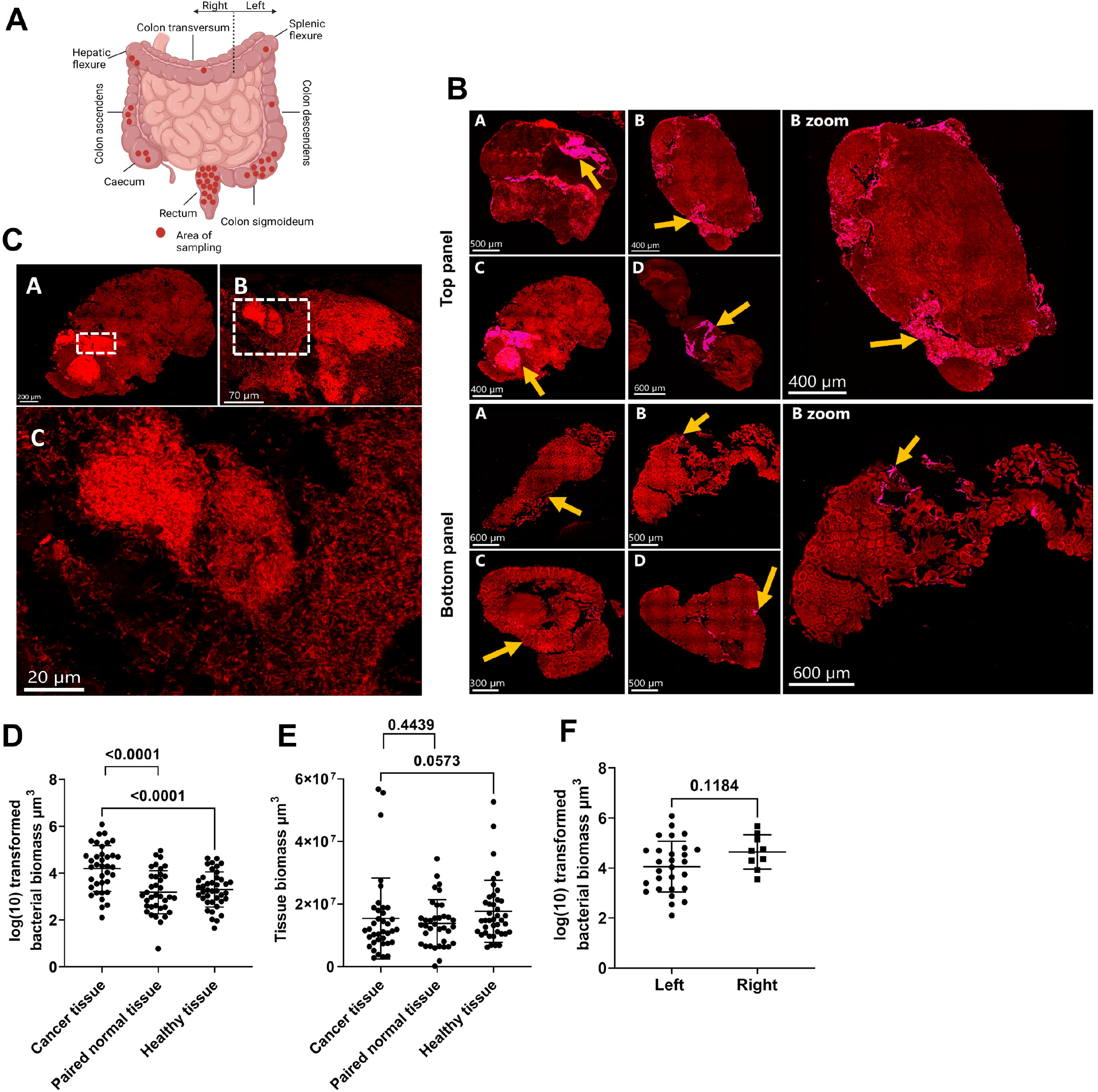
Assessment of bacterial biomass in biopsies collected from patients with and without CRC. **A)** Diagram showing the anatomical sampling of CRC biopsies. **B)** Cross-sections of tumour biopsies (top panel) and healthy tissue biopsies (bottom panel) showing the distribution of bacterial biomass. Yellow arrows indicate the area with bacterial biomass and a mask was created to overlay areas with bacteria with the Imaris software through thresholding of fluorescence intensity. Tissue was visible via autofluorescence. Scale bars are shown in the lower-left corner of all images. **C)** Morphological evidence of bacteria in the cross-section of a tumour biopsy from the top-panel (image C) with encircled areas in A and B representing the magnification in B and C, respectively. **D)** Logarithmic (Log) transformed bacterial biomass measured in cubic micrometers (µm^3^) from collected biopsies. **E)** Total tissue biomass (bacteria and tissue) measured in µm^3^ from collected biopsies. **F)** Log-transformed bacterial biomass on the left- and right-sided tumour biopsies measured in µm^3^. Statistical comparison was carried out with paired and unpaired t-tests **(D)**, paired t-test and Mann-Whitney test **(E)**, and unpaired t-test **(F)**. Bars represent standard deviation (SD). A p-value ≤ 0.05 was considered statistically significant.

**Table 1.**
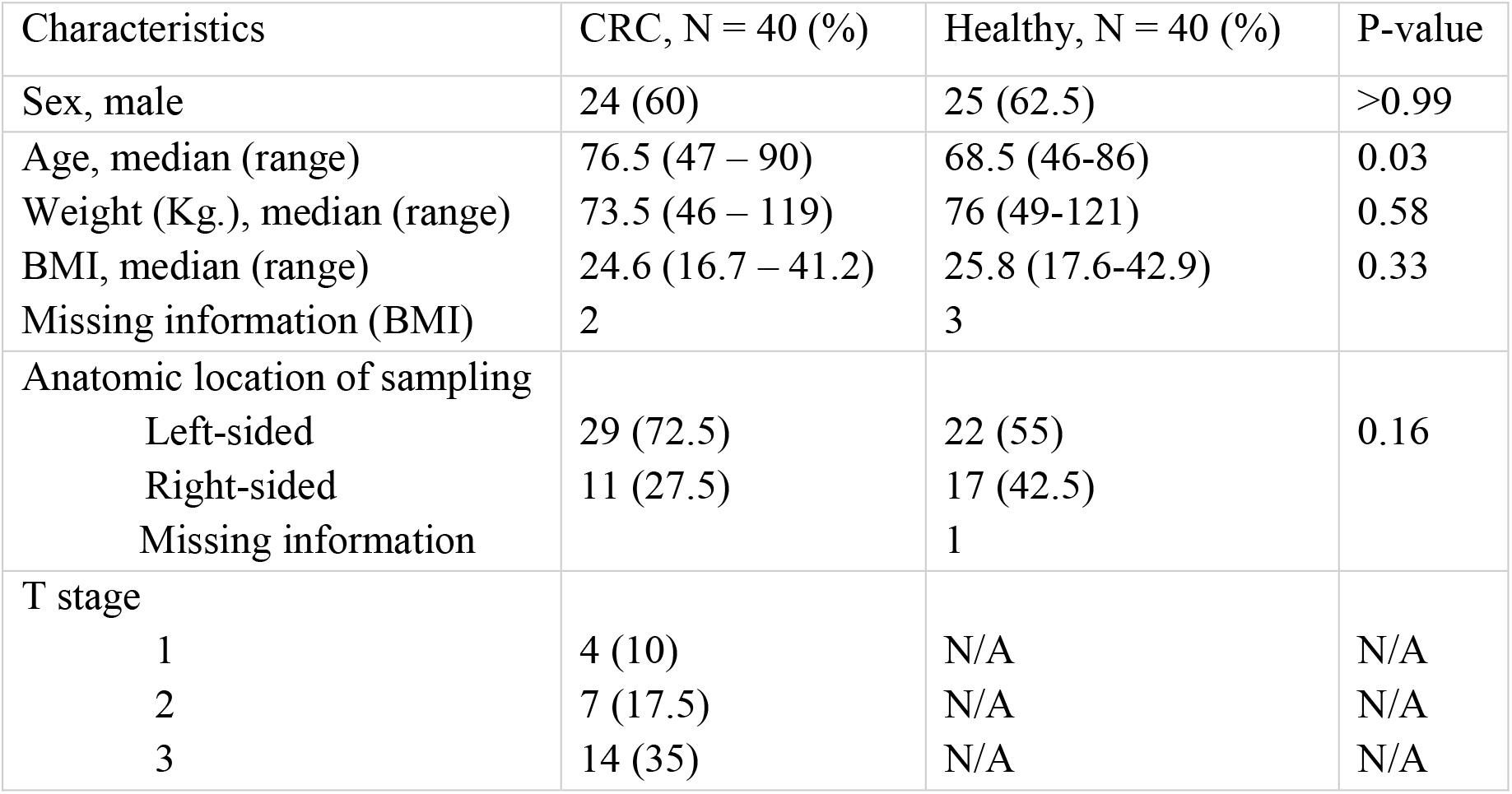
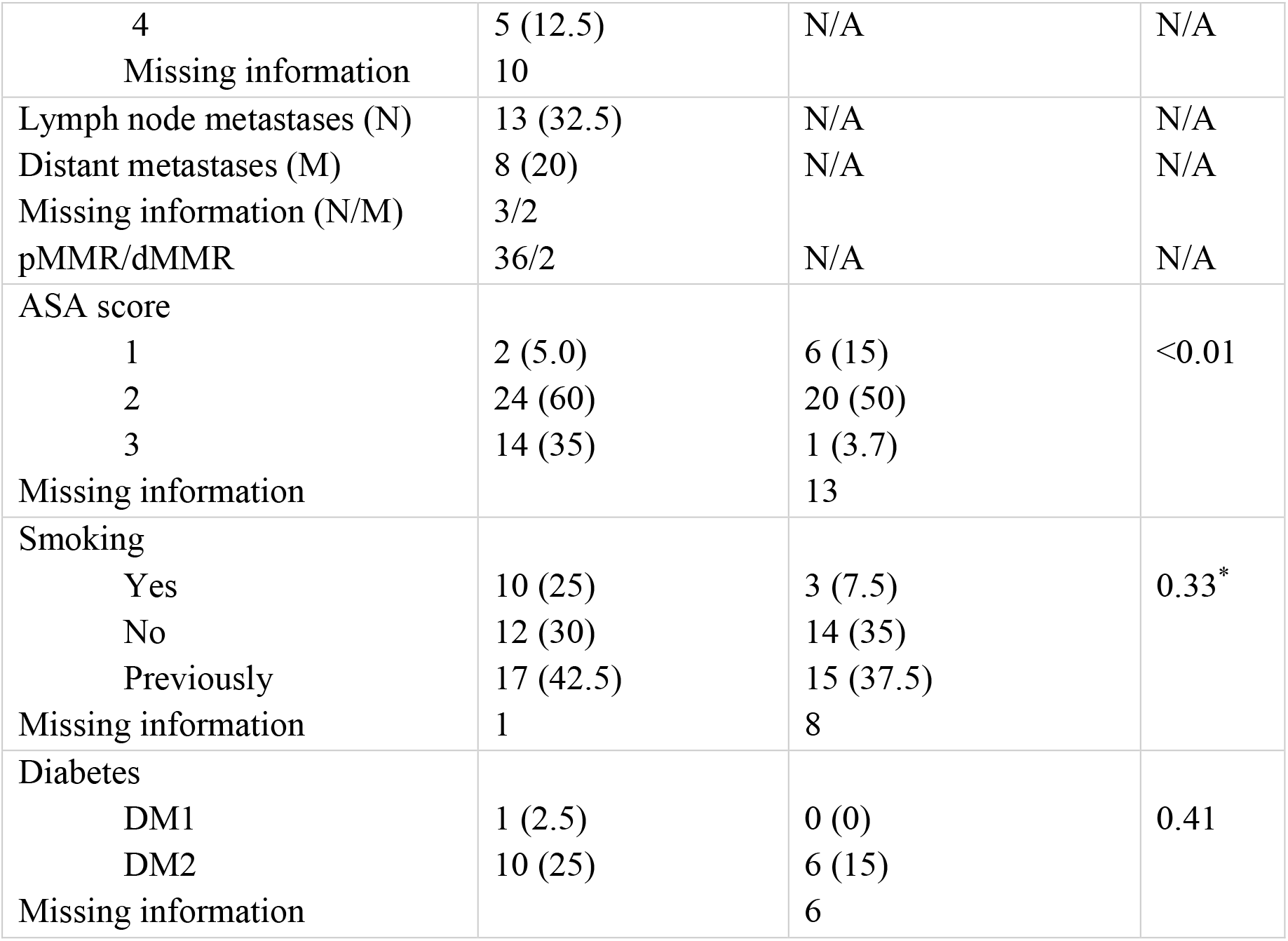
Characteristics of included patients with CRC and healthy subjects. ASA, American Society of Anesthesiologists. BMI, Body Mass Index (Kg/m^2). DM, Diabetes mellitus. *Current and previous smoking has been pooled for statistical analysis. Continuous data was tested with a two-sided student t-test or Mann-Whitney Test, and categorical data was tested with a chi-square test. A p-value ≤ 0.05 was considered statistically significant.

### Increased bacterial biomass was observed in CRC tissue

Bacterial biomass (µm^3^) was quantified systematically in biopsies using panbacterial PNA-FISH. Three pairs of samples (CRC and paired normal tissue) were excluded due to non-cancerous origin (n=2) and incorrect processing (n=1). In one case, it was not possible to sample paired normal tissue. Thus, 113 mucosal biopsies from 37 patients with CRC and 40 healthy persons were examined. No differences were observed between groups after removing samples (Table S1). The bacterial biomass displayed a tissue-invasive phenotype in CRC biopsies, while bacteria were generally localized along the epithelial lining of healthy colon biopsies (representative images shown in Figures 1B and 1C). Large patches of aggregated bacteria (biofilm) were frequently observed in CRC tissue, and bacterial biomass was higher in CRC tissue compared to paired normal and healthy tissue (Figure 1D). The mean bacterial biomass was 11 to 17-fold higher in CRC tissue (0.70% of the total biomass) compared to paired normal (0.06%) and healthy tissue (0.04%). There were no differences in biopsy sizes across the groups (Figure 1E). When stratifying the bacterial biomass into the respective anatomic locations of sampling (Figure 1A), a stepwise increase in mean bacterial biomass was observed from the rectum (3.93 ± 0.90 SD) over sigmoideum (4.32 ± 1.01 SD) to colon ascendens (4.35 ± 0.65 SD) and caecum (4.76 ± 0.72 SD); however, this was not significant. Similarly, no differences (mean difference = 0.59 ± 0.37 SEM, p=0.12) were observed when stratifying into left- and right-sided tumours (Figure 1F), as previously reported ^38^. Bacterial biomass was not associated with tumour staging (T1-T4), lymph node metastasis (N), or distant metastasis (M) (Figure S2).

### The prevalence of *Fusobacterium spp.* correlated with increased bacterial biomass and virulence expression profile in CRC tissue

Species-specific PNA-FISH was used to assess the prevalence and contribution of *Fusobacterium spp.* and *B. fragilis* to bacterial biomass in CRC. Successive separation of the probes was initially tested in spiked tissue and tumour tissue (Figure 2A). *Fusobacterium spp.* were observed in 24 out of 37 (64.9 %) tumour biopsies, 18 out of 36 (50.0 %) paired normal biopsies, and 14 out of 40 (35.0 %) healthy biopsies*. B. fragilis* was observed in 19 out of 37 tumour biopsies (51.4 %), 15 out of 36 paired normal biopsies (41.7 %), and 13 out of 40 healthy biopsies (32.5 %). The prevalence of *Fusobacterium spp.* was significantly higher in CRC tissue than in healthy tissue (Figure 2B). No difference in prevalence was observed between groups for *B. fragilis*. Interestingly, a higher prevalence of *B. fragilis* (Figure 2C) and *Fusobacterium spp.* (Figure 2D) was observed in right-sided tumours, suggesting anatomical preference. The prevalence of *Fusobacterium spp.* and *B. fragilis* was not associated with tumour staging (T1-T4), lymph node metastasis (N), or distant metastasis (M) (Figure S2). Microscopy revealed that *Fusoacterium spp.* formed a substantial proportion of the mixed-species biofilms in CRC tissue (representative image in Figure 2E), suggesting superior adhesion or facilitated co-adhesion of other bacteria. Adhesion of *Fusobacterium spp.* to the epithelial cells or other bacteria through its virulence factors is well described ^39–42^. A sub-group analysis revealed that samples positive with *Fusobacterium spp.* had a higher mean percentage of bacterial biomass (1.06 % vs. 0.14 %) than those without *Fusobacterium spp.* (Figure 2F).

**Figure 2.**
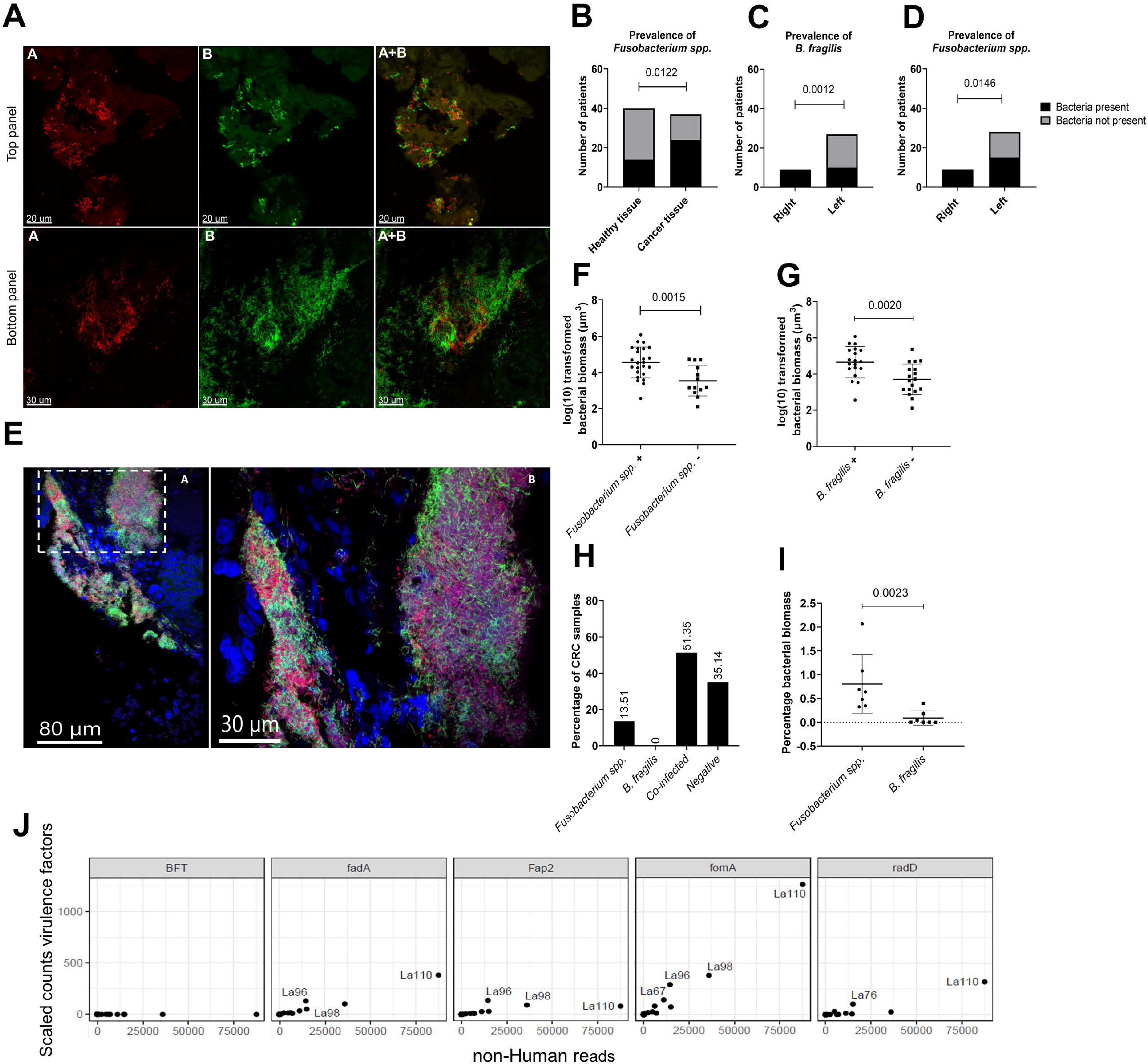
Prevalence of *Fusobacterium spp*. and *B. fragilis* in CRC, paired normal, and healthy tissue. **A)** Representative images showing the qualitative separation of FUS714 (*Fusobacterium spp.*) from Bfrag-998 (*B. fragilis*) in spiked tissue (top panel) and tumour tissue (bottom panel). **B)** Prevalence of *Fusobacterium spp.* in healthy tissue and cancer tissue. **C+D)** Prevalence of *B. fragilis* (C) and *Fusobacterium spp.* (D) in left- and right-sided tumours. **E**) Representative images showing mixed-species biofilms with *B. fragilis* (red), *Fusobacterium spp.* (green), other bacteria (purple), and host cells (blue). Image A is an overview image, and the white bracketed box shows the enlarged area in image B. **F+G)** Correlation between the prevalence of *Fusobacterium spp.* (B) and *B. fragilis* (C) and logarithmic (Log) transformed bacterial biomass in CRC tissue measured in cubic micrometers (µm^3^). **H)** Percentage of CRC samples positive with either *Fusobacterium spp.*, *B. fragilis*, co-infected or negative. **I)** The percentage of bacterial biomass for *B. fragilis* and *Fusobacterium spp.* on a subset of co-infected samples (n=7) with high bacterial biomass. **J**) Scaled counts assigned to the *B. fragilis* toxin (BFT) and *F. nucleatum* virulence factors FadA, Fap2, FomA, and RadD. Scale bars are shown in the lower-left corner of all images (A+E). Statistical comparison was carried out with Fisher’s exact tests (**B+C+D)**, unpaired t-tests (**F+G),** and Mann-Whitney tests (**I)**. Bars represent standard deviation (SD). A p-value ≤ 0.05 was considered statistically significant.

Similarly, the mean bacterial biomass in samples with *B. fragilis* (Figure 2G) was higher than those without; however, we hypothesized these findings were confounded due to co-infection by *Fusobacterium ssp.* (Figure 2H). Therefore, the specific bacterial biomass of *Fusobacterium spp.* and *B. fragilis* was analyzed in seven co-infected samples with high bacterial biomass. *Fusobacterium spp.* was more abundant than *B. fragilis* in these samples, and a significant difference was observed in the mean percentage of bacterial biomass (Figure 2I). Moreover, the expression patterns of *B. fragilis* enterotoxin (BFT) and the *F. nucelatum* virulence factors FadA, Fap2, FomA, and radD were analyzed. Only the virulence factors of *F. nucleatum* were enriched in CRC samples (Figure 2J), showing that no active enterotoxin-producing *B. fragilis* were present and that *Fusobacterium ssp*. expressed the virulence factors necessitated for epithelial adherence and co-adherence to other bacteria.

### Bacterial biomass was associated with acute inflammation and co-localized with necrotic areas in CRC tissue

Two blinded pathologists scored CRC biopsies to determine if bacteria influenced the TME. Inflammation scores, reflecting PMNs and lymphocyte infiltration, were given for acute and chronic inflammation^24 25^. Samples with high bacterial biomass had a higher degree of acute inflammation than samples with low bacterial biomass (Figure 3A). Similarly, there was a higher degree of inflammation in samples with *B. fragilis* and *Fusobacterium spp.*; however, this was not significant (Figures 3B and 3C). No inflammation was observed in healthy biopsies (data not shown). Moreover, in 18 samples (85.71 %) with high bacterial biomass (n = 21), it was observed that bacteria and necrotic tissue were co-localized (representative images shown in Figure 3D), suggesting an anatomical preference for biofilm growth or that bacteria are involved in the malignant transformation.

**Figure 3.**
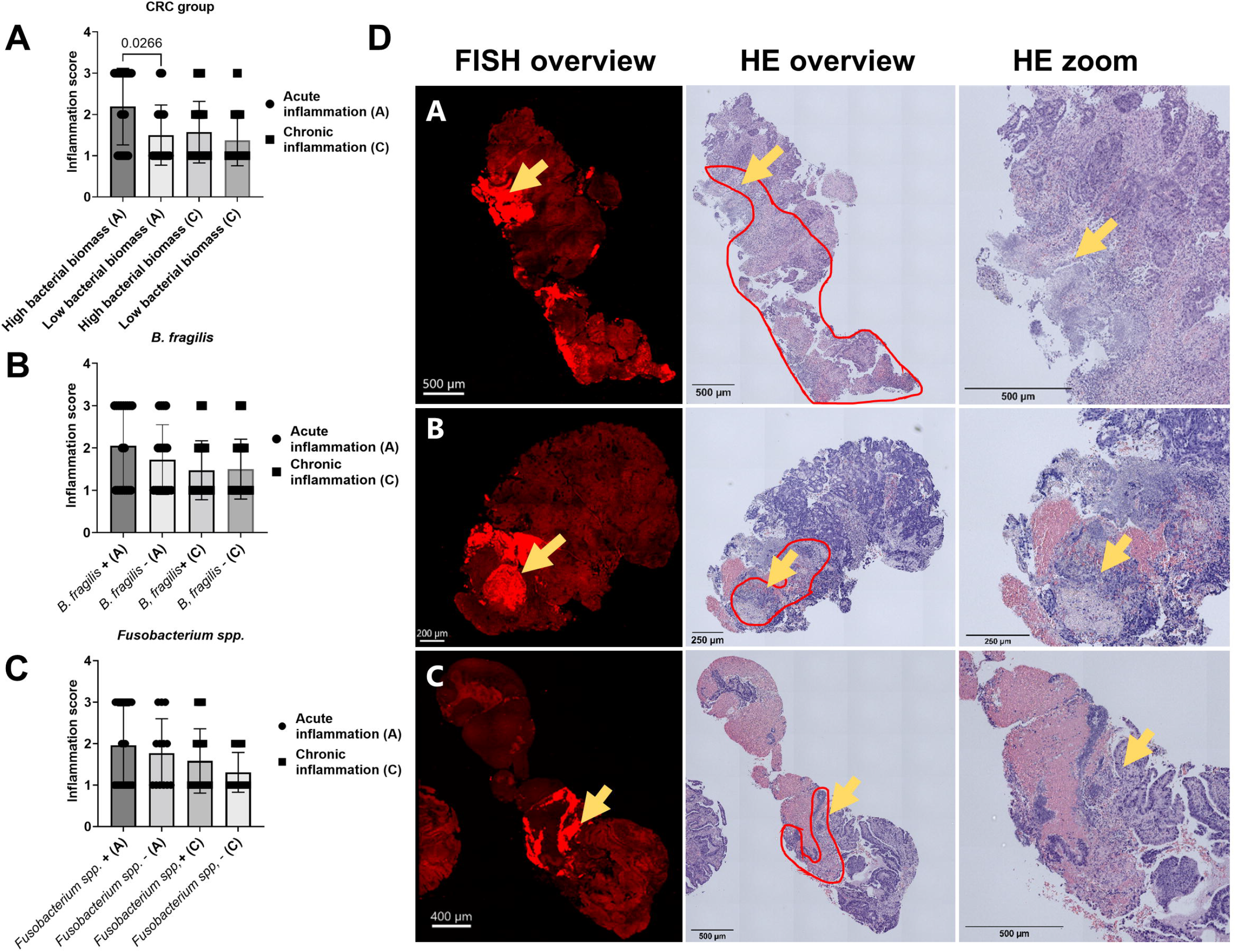
Histopathological evaluation of the bacterial influence on the TME. **A)** Inflammation score for CRC samples with high and low bacterial biomass, divided into acute inflammation and chronic inflammation. **B**+**C**) Inflammation score for CRC samples with (+) and without (−) *B. fragilis* (B) and *Fusobacterium spp.* (C), divided into acute and chronic inflammation. **D**) Representative images showing the co-localization of bacterial biomass and necrotic tissue on cross-sections of tumour biopsies. The red encircled area indicates the area with necrosis, and the yellow arrows indicate the area with bacterial biomass. All biomass measurements were performed with the Imaris software through thresholding of fluorescence intensity. Tissue was visible via autofluorescence. Scale bars are shown in the lower-left corner of all images. Statistical comparison was carried out with Mann-Whitney tests (A+B+C). Bars represent standard deviation (SD). A p-value ≤ 0.05 was considered statistically significant.

### Higher counts for bacteria, *F. nucleatum,* and *B. fragilis* were observed in CRC tissue

RNA sequencing was performed on 118 samples to assess the bacterial activity within CRC, paired normal, and healthy tissue. Two samples were excluded: one healthy sample due to a failed library preparation and one paired normal sample because it was not possible to collect tissue. RNA-seq reads were taxonomically assigned, classified, scaled, and quantified for each sample. In all samples, the highest counts were assigned to Eukaryota (mainly human), bacteria, viruses, and archaea in the order mentioned (Figure S1). Bacterial counts were in the order of 1:100 compared to human counts. Sample type described the majority of sample variation (Figure S3). Bacterial counts were higher in CRC tissue (Figure 4A) when compared to paired normal (p<0.003, Wilcoxon signed-rank test) and healthy tissue (p=0.03, Wilcoxon rank-sum-test). The groups did not differ in alpha diversity (Figure 4B). The Fusobacteria phylum (Figure 4C) showed higher counts in CRC tissue than in healthy (p<0.001, Wilcoxon rank-sum-test) or paired normal tissue (p<0.001, Wilcoxon signed-rank test). In general, Firmicutes and Proteobacteria were the dominant phyla across all groups (Figure 4D–4F), while an increase of Bacteroidota, Fusobacteria, and Actinobacteria was mainly observed in the CRC group (Figure 4G–4I). Similar to previous findings, *F. nucleatum* and *B. fragilis* counts were higher in CRC tissue (Figure 4J) compared to healthy (p<0.001, Wilcoxon rank-sum-test) and paired normal tissue (p<0.001, Wilcoxon signed-rank test). Nine and seven samples deviated from the normal count distribution assigned to *F. nucleatum* and *B. fragilis*, respectively (Figure S1). In the nine samples, counts assigned to *F. nucleatum* comprised approximately 79% of the total counts assigned to the Fusobacteria phylum. Other *Fusobacterium spp.* were present in CRC samples; however, *F. nucleatum* was most abundant (Figure 2K). These results suggest that increased findings of *Fusobacterium ssp.* in the samples subjected to microscopy probably were due to the presence of *F. nucleatum*.

**Figure 4.**
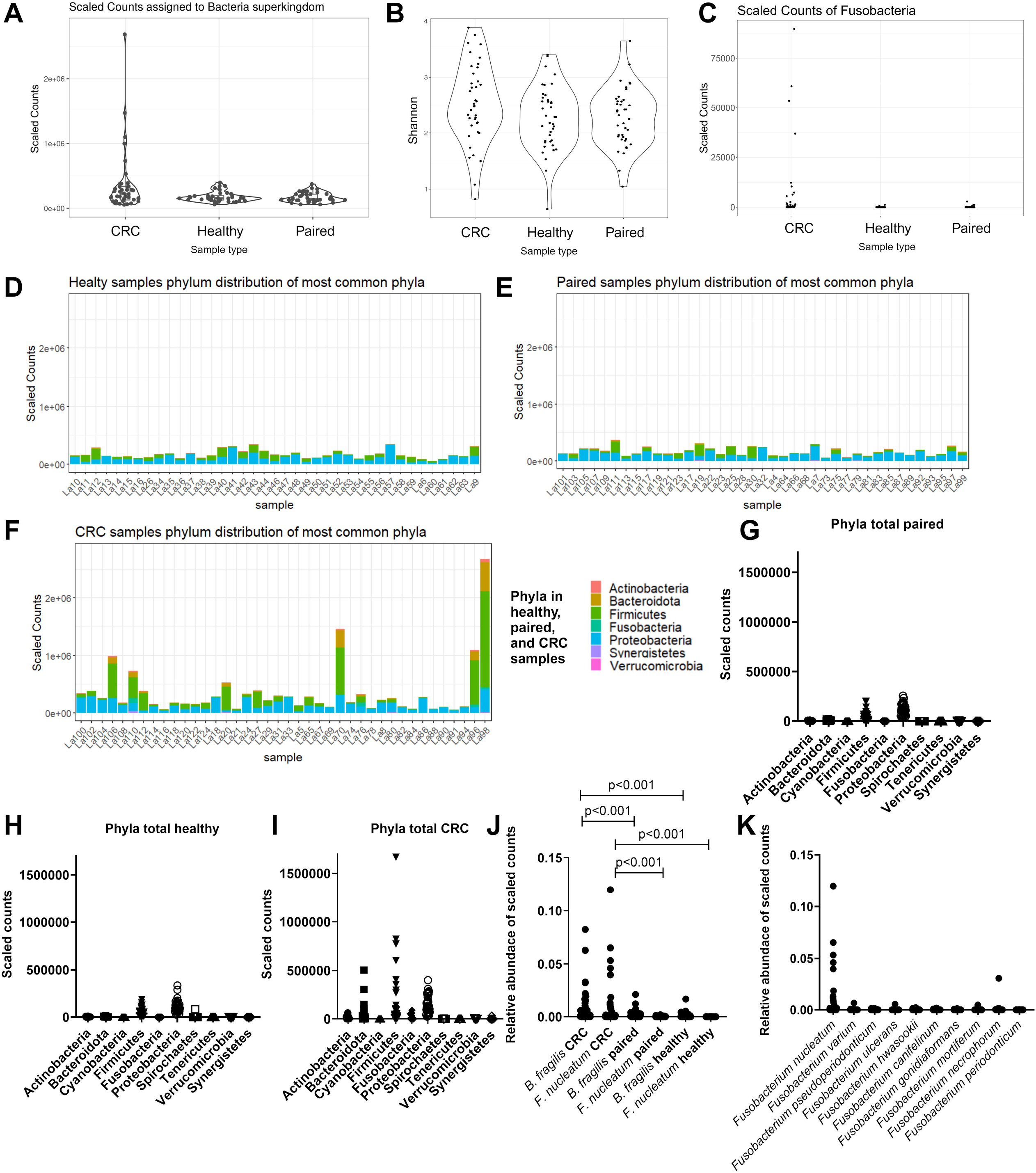
Characterisation of bacterial richness and diversity in CRC, paired normal, and healthy tissue. **A)** Scaled counts assigned to bacteria in CRC, healthy, and paired normal tissue. **B)** The alpha diversity in CRC, healthy, and paired normal tissue was compared with the Shannon index. **C)** Scaled counts assigned to Fusobacteria in CRC, healthy, and paired normal tissue. **D+E+F)** Scaled counts were assigned to the seven most dominant phyla for each sample in paired normal (D), healthy (E), and CRC tissue (F), respectively. **G+H+I)** Scaled counts were assigned to the ten most dominant phyla across all samples in paired normal (G), healthy (H), and CRC tissue (I), respectively. **J)** Relative abundance of scaled counts assigned to *B. fragilis* and *F. nucleatum* in CRC, paired normal, and healthy tissue. **K)** Relative abundance of scaled counts assigned to *Fusobacterieum spp*. in CRC tissue. Statistical comparison was carried out with Wilcoxon signed-rank test and Wilcoxon rank-sum-test (**A+C+J**), paired and unpaired t-tests (**B**), and Mann-Whitney tests (I+J+K). Bars represent standard deviation (SD). A p-value ≤ 0.05 was considered statistically significant.

### Higher bacterial activity in CRC tissue affects host transcription and immune phenotype

Six CRC samples containing elevated levels of bacterial RNA were identified in our cohort (Figure S1C). To assess the effect of increased bacterial activity on the TME, differential gene expression and functional enrichment analyses were performed between the CRC samples containing high (n=6) and low (n=34) bacterial RNA. This analysis identified 332 significantly differentially expressed genes due to bacterial activity in CRC, where 252 showed increased expression with increased bacterial activity (Figure 5A). These included increased expression of several proinflammatory cytokines (*CXCL6*, *CXCL8*, *CXCL9*, *IL1B*, *IL6*, *CCL3*, and *CCL7*), defensins (*DEFA1*, *DEFA3*, *DEFA4*, *DEFB125*, *DEFB129*, and *DEFB131*), matrix-metalloproteases (*MMP1*, *MMP 12*, and *MMP13*) and other immunomodulatory factors and receptors (*S100A8*, *MEFV*, *CD86*, *CSF3*, *FPR1*, *PTGS2*, and *TLR*). These genes represented significant enrichment of IL-10 signalling, defensin, chemokine, and other pathways (Figure 5B). Further, metabolic pathways involving UDP-glucuronosyltransferases (*UGT1A1, UGT1A10, UGT1A4, UGT1A7, UGT1A8, UGT1A9*, *UGT2B15*, and *UGT2B17*), alcohol dehydrogenases (*ADH1B* and *ADH1C*), and cytochrome P450 genes (*CYP2B6, CYP2C18, CYP2C19, CYP2B6, CYP2C18, CYP2C19,* and *CYP4F12*) showed significantly increased expression in samples with low bacterial activity. Interestingly, many of these significantly enriched pathways in CRC samples with low bacterial activity (Figure 5B) overlapped with significantly enriched pathways in healthy samples compared with CRC samples (Figure S3). We then investigated whether an increased total bacterial or species-specific activity of *F. nucleatum* and *B. fragilis* affected the TME immune phenotype in CRC tissue. This analysis integrated several existing cell deconvolution and immune scoring systems to develop a more robust estimate and was grouped by whether the output was a fraction (Figure 5C) or a normalized score (Figure 5D). This analysis included 115 samples (3 samples were identified as outliers and removed from the dataset), and 26.920 genes were used. Four clusters were identified separating CRC and healthy samples (paired normal and healthy tissue) with high and low immune scores (Figure 5E). The overall bacterial and species-specific activity showed associations with clusters but not with specific immune-cell subtype abundance herein.

**Figure 5.**
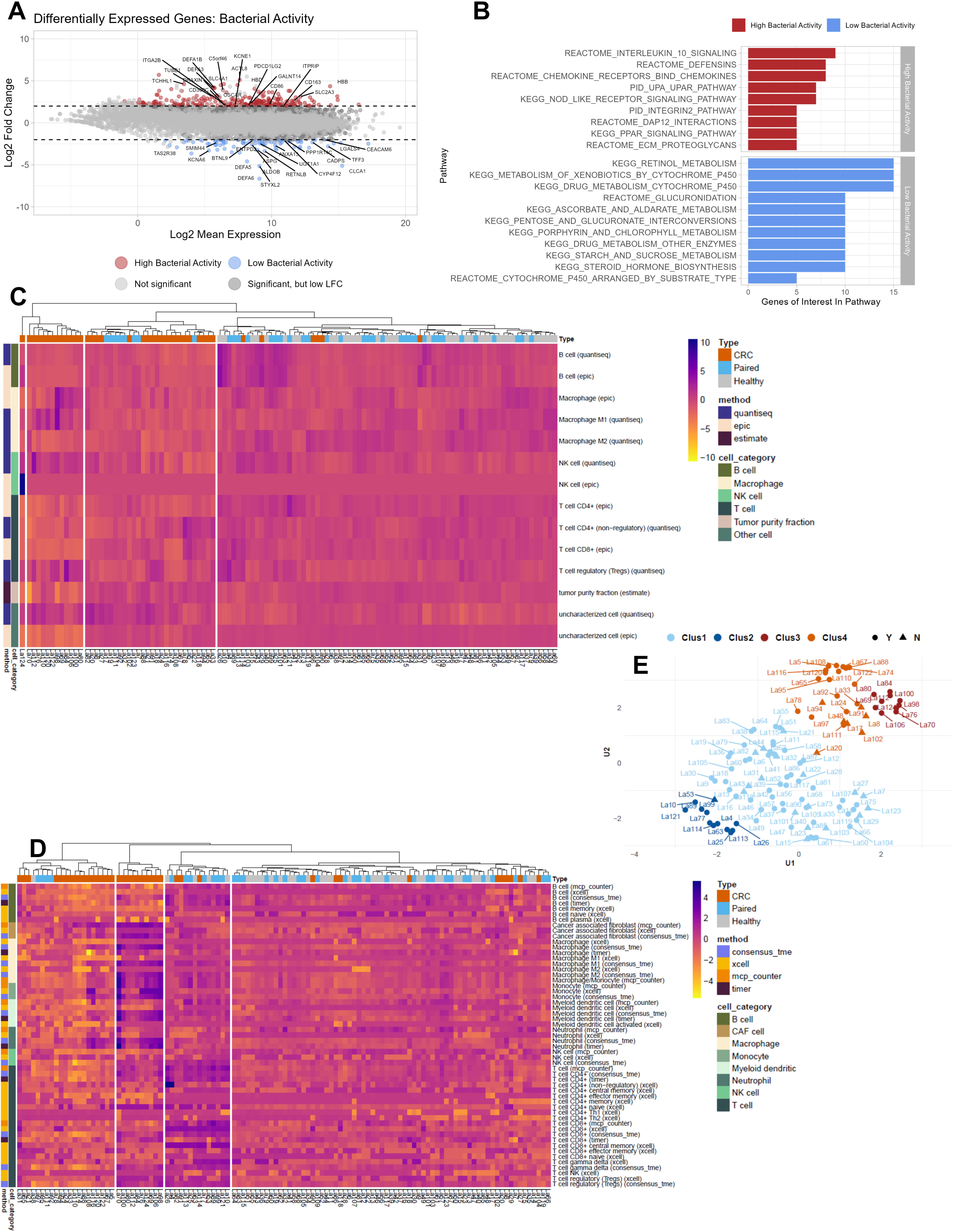
Differentially expressed genes (DEGs) and tissue immune phenotype. **A)** MA plot showing the distribution of significantly differentially expressed genes between CRC tissue with high and low bacterial activity. Coloring highlights the 20 most significant DEGs with an adjusted p-value less than 0.05 and absolute log2 fold-change >2. **B)** The Kyoto Encyclopedia of Genes and Genomes (KEGG), Pathway Interaction Database (PID), and REACTOME (a database of reactions, pathways, and biological processes) databases were used to identify pathways that were significantly enriched or decreased in CRC tissue with high and low bacterial activity. **C+D)** Heatmaps showing immune cell profiles in CRC, healthy, and paired normal tissue, presented as fractions from the quantisec, epic, and estimate immune scoring systems (C) and normalized scores from the concensus_tme, xcell, mcp_counter, and timer immune scoring systems (D). Coloring from yellow (−4) to purple (4) indicates the degree of infiltration, where purple is high infiltration. **E)** Four immune cell profile clusters were defined (Clus1-Clus4) in this study based on the hierarchical clustering of samples according to immune cell infiltration in the heatmaps. Y and N indicate whether samples were stable to the assigned clusters.

However, when differentiating between individual immune cell subtypes across sample types (CRC, paired normal, and healthy tissue), bacterial counts affected the infiltration of macrophage/monocyte, myeloid dendritic cells, regulatory T-cells, tumour purity fraction, and tumour purity score (Table 2). *F. nucleatum* did not correlate with the infiltration of any immune cells across sample type, while *B. fragilis* correlated with the infiltration of neutrophils and the tumour purity fraction score (Table 2). The analysis was also conducted excluding paired normal samples, as dysbiosis has been suggested to occur in the whole colon ^43^. Similar hierarchical sample clustering was observed (Figures S5A and S5B). Only species-specific activity was associated with clusters but not specific immune-cell subtype abundance. When assessing the influence on sample types, *F. nucleatum* correlated with the infiltration of effector memory CD4+ T-cells, while *B. fragilis* did not correlate with the infiltration of any immune cell subtypes. Of particular interest, the list of immune cell subtypes was extended for total bacterial counts (Table 2), including the immune and microenvironment scores. An overview of all the immune cell subtypes and methods used can be found in Tables S2-S7.

**Table 2.** Bacterial groups impacting infiltration of specific immune cells in CRC tissue.

**Paired normal and healthy tissue samples were used as controls.**
| **Specific counts** | **Affected immune cell group** | **Strength of association** | **Methods finding an effect** |
| --- | --- | --- | --- |
| Bacteria | Macrophage/Monocyte | $p < 0.01$ | 1 out of 1 |
| Bacteria | Myeloid dendritic cells (activated) | $p < 0.05$ | 1 out of 1 |
| Bacteria | Regulatory T-cells | $p < 0.001$ | 1 out of 2 |
| Bacteria | Tumour purity fraction | $p < 0.001$ | 1 out of 1 |
| Bacteria | Tumour purity score | $p < 0.05$ | 1 out of 1 |
| *B. fragilis* | Neutrophils | $p < 0.05$ | 1 out of 4 |
| *B. fragilis* | Tumour purity fraction | $p < 0.05$ | 1 out of 1 |

**Healthy tissue samples were used as controls.**
| Specific counts | Affected immune cell group | Strength of association | Methods finding an effect |
| --- | --- | --- | --- |
| Bacteria | Eosinophils | $p < 0.05$ | 1 out of 2 |
| Bacteria | Hematopoietic stem cell | $p < 0.01$ | 1 out of 1 |
| Bacteria | Immune score | $p < 0.05$ and $p < 0.01$ | 2 out of 3 |
| Bacteria | Macrophage M2 | $p < 0.05$ and $p < 0.05$ | 2 out of 2 |
| Bacteria | Macrophage/Monocyte | $p < 0.01$ | 1 out of 1 |
| Bacteria | Microenvironment score | $p < 0.05$ | 1 out of 1 |
| Bacteria | Monocyte | $p < 0.05$ and $p < 0.01$ | 2 out of 3 |
| Bacteria | Myeloid dendritic cells (activated) | $p < 0.001$ | 1 out of 1 |
| Bacteria | Regulatory T-cells | $p < 0.001$ | 1 out of 2 |
| Bacteria | Tumour purity fraction | $p < 0.0001$ | 1 out of 1 |
| Bacteria | Tumour purity score | $p < 0.01$ | 1 out of 1 |
| *F. nucleatum* | T cell CD4+ effector memory | $p < 0.05$ | 1 out of 1 |

## DISCUSSION

### Bio-geography and bacterial biomass in CRC samples

A tissue-invasive phenotype was observed in tumour biopsies with significant enrichment of bacterial biomass compared to paired normal and healthy tissues. These results are in accordance with previous findings, where approximately 50% of CRC samples and 13% of healthy samples harboured bacterial biofilms, with a higher density of bacteria in CRC samples ^38^. In addition, when stratifying data into anatomic compartments of sampling, right-sided tumours from three different cohorts have shown invasive biofilms in 93 % of the cases, whereas it was observed in 27 % of cases for left-sided tumours ^44^. These findings support the findings in this study, where a trend towards higher bacterial biomass was observed in right-sided tumours. In agreement with previous studies, bacterial biomass did not correlate with tumour stage, lymph node metastasis, or distant metastasis (Figure S2) ^38^ ^44^. Interestingly, there was a correlation between high bacterial biomass and the degree of infiltrating PMNs in CRC samples, meaning biofilms can alter the TME and provoke an inflammatory response. *F. nucleatum* has previously been observed in ulcerated regions ^45^, aligning with our results, where bacterial biomass co-localized with necrotic tissue, further implicating biofilms in the inflammatory response.

### Core pathogens in CRC tissue

Consistent with the microscopic findings, increased bacterial richness or transcriptional activity was observed in CRC tissue compared to healthy and paired normal tissue. Mucosal biopsies were enriched with Proteobacteria and Firmicutes across all groups, with the highest counts assigned to CRC tissue. In addition, there was an increase in counts assigned to Bacteroidota, Fusobacteria, and Actinobacteria in the CRC group. A recent study by Zhao et al. described the consensus mucosal microbiome from 924 tumours, including eight RNA datasets across different geographical locations, and found the same phyla elevated in CRC tissue with no difference in alpha diversity, similar to our findings ^46^.

*F. nucleatum* has been well studied over the last couple of years, and multiple studies have found an enrichment of this bacterium in CRC tissue ^44^ ^47^ ^48^. In our study, *F. nucleatum* was significantly enriched in CRC tissue and was the dominant species in the Fusobacteria phylum. Interestingly, *Fusobacterium spp.* was more prevalent in right-sided tumours. A previous study could not find a correlation between *F. nucleatum* and right-sided tumours ^44^ however, they employed sequencing techniques specific to *F. nucleatum,* whereas a probe targetting *Fusobacterium spp.* was used in this study, which could explain the difference. In support of this notion, a study by Tahara *et* al. found a correlation between the enrichment of *Fusobacterium spp.* and subsets of colorectal cancers known to dominate in right-sided tumours ^49^. Of particular interest, we found that *Fusobacterium spp.* was associated with higher bacterial biomass, suggesting superior adhesion of *Fusobacterium spp.* to the host tissue or that *Fusobacterium spp.* facilitated the co-adhesion of other bacteria, as reported for *F. nucleatum* in periodontal diseases ^39^ ^50^. *F. nucleatum* has tissue-adhesive and co-aggregating properties qua its virulence factors FadA Fap2, RadD, and FomA ^39–42^. Accordingly, FadA, Fap2, RadD, and FomA expression were detected in CRC samples with high *F. nucleatum* counts. To our knowledge, this is the first study to show that *Fusobacterium spp.* are associated with increased bacterial biomass in CRC. A recent study employing laser-microdissection 16S rRNA gene sequencing on tissue samples also found enrichment of *B. fragilis* in right-sided tumours ^51^. Similarly, we found that *B. fragilis* was enriched on the right side of the colon. *B. fragilis* displays high strain diversity in the human gut and can be divided into toxigenic (BFT-producing) or non-toxigenic strains, both implicated in CRC tumorigenesis ^52^. We could not detect the expression of BFT in our study, suggesting that no active toxin-producing *B. fragilis* were present. *B. fragilis* toxins are more common in right-sided tumours ^53^, and the inclusion of a few right-sided tumour biopsies in our study might influence these findings. In contrast to microscopy findings, a higher enrichment of *B. fragilis* was observed in CRC samples compared to healthy and paired normal tissue, emphasizing the importance of using complementary methods to characterize the mucosa-associated microbiota.

### Bacterial activity and host-associated transcriptional and immunologic responses

Samples submitted to RNA sequencing varied in terms of bacterial activity and the expression profile of genes. Samples with high bacterial activity exhibited a pro-inflammatory signature with increased expression of genes coding for pro-inflammatory cytokines, defensins, matrix-metalloproteases, and other immunomodulatory factors. These findings contrasted with the expression profile in samples with a low bacterial activity where the enriched pathways overlapped with significantly enriched pathways in healthy samples. These findings highlight that increased bacterial activity negatively impacts the local TME in terms of an increased inflammatory response, which can fuel or sustain a pro-tumourigenic environment.

In line with this, total bacterial counts affected several immune cell subpopulations in CRC tissue, including the overall immune and microenvironment scores. The species-specific association was modest and included associations between *B. fragilis* and neutrophil infiltration and between *F. nucleatum* and effector memory CD4+ T-cell infiltration. Previous studies have evaluated the effect of *F. nucleatum* on CD4+ T-cell activity, with conflicting findings ^11^ ^12^ ^54^; however, to our knowledge, this is the first time that *B. fragilis* has been associated with the infiltration of neutrophils. These results indicate that the species-specific contribution to immune cell infiltration only constitutes a part of the immunologic response and emphasize the importance of widening the bacterial scope when investigating the bacterial role in CRC carcinogenesis.

## CONCLUSION

CRC core pathogens such as *F. nucleatum* and *B. fragilis* are highly prevalent in CRC tissue, specifically right-sided tumours. *F. nucleatum* plays a role in the build-up of mixed-species biofilms, possibly due to the expression of tissue-adhesive and co-aggregating virulence factors, resulting in an increased accumulation of bacterial biomass and higher inflammatory response. These findings suggest a reduction in the bacterial biomass as a potential target to reduce inflammation-driven CRC carcinogenesis; however, there is a lack of clinical studies in this area, and future studies should explore the clinical implications of reducing bacterial biomass or species-specific antimicrobial targeting of *Fusobacterium spp.* While *F. nucleatum* and *B. fragilis* were enriched in CRC tissue, their effect on the TME and infiltration of immune cells was modest. In contrast, the collective presence of bacteria seemed more relevant in altering the immune phenotype and regulating genes and critical pathways. These observations confirm the narrative of the involvement of *F. nucleatum* and *B. fragilis* in CRC carcinogenesis while highlighting the importance of also widening the bacterial scope beyond CRC core pathogens when deciphering the role of bacteria in CRC carcinogenesis.

## ACKNOWLEDGEMENTS

We want to thank Veronica Drejer for her sublime help with RNA purification and preparation of samples for RNA sequencing. We would also like to thank Britt Capellen from the Center of Surgical Science for the initial sample preparation prior to RNA purification. The illustration of the sampling area in Figure 1 was created with Biorender.com.

## AUTHOR CONTRIBUTIONS

Conceptualization, L.K., I.G., and T.B.; methodology, L.K., B.G.F, I.G., and T.B.; investigation, L.K., B.G.F., M.R.J., and A.G.G.; data curation, L.K., B.G.F., H.Z., T.B.T., and K.H-R.; data analysis, L.K. B.G.F., T.B.T., and H.Z.; generation of figures, L.K. B.G.F., T.B.T., and H.Z.; writing – original draft, L.K.; writing – review & editing, All authors.; supervision, H.R, I.G. and T.B.; project administration, L.K.; funding acquisition, I.G., and T.B.

## SUPPLEMENTAL INFORMATION LEGENDS

**Table S1 – Characteristics of included patients with CRC and healthy persons.** ASA, American Society of Anesthesiologists. BMI, Body Mass Index (Kg/m^2). DM, Diabetes mellitus. *Current and previous smoking has been pooled for statistical analysis. Continuous data were tested with a two-sided student t-test or Mann-Whitney test, and categorical data were tested with a chi-square test. A p-value ≤ 0.05 was considered statistically significant.

**Table S2 – Bacterial counts impacting specific immune cells across the seven methods that score immune cell infiltration.** NA = Not applicable because the method does not report that type of immune cell. ns = not significant. * = p<0.05, ** = p<0.01, *** p<0.001, **** p<0.0001. Statistical comparison was carried out with Ordinary Least Squares regression to determine which independent variables (sample type, bacterial count, read count) explain the dependent outcome variable (the immune score). Highlighted rows indicate immune cell sub-populations only affected in CRC tissue by bacterial counts. A p-value ≤ 0.05 was considered statistically significant.

**Table S3 – *Bacteroides fragilis* impacting specific immune cells across the seven methods that score immune cell infiltration.** NA = Not applicable because the method does not report that type of immune cell. ns = not significant. * = p<0.05. Statistical comparison was carried out with Ordinary Least Squares regression to determine which independent variables (sample type, bacterial count, read count) explain the dependent outcome variable (the immune score). Highlighted rows indicate immune cell sub-populations only affected in CRC tissue by bacterial counts. A p-value ≤ 0.05 was considered statistically significant.

**Table S4 – *Fusobacterium nucleatum* impacting specific immune cells across the seven methods that score immune cell infiltration.** NA = Not applicable because the method does not report that type of immune cell. ns = not significant. Statistical comparison was carried out with Ordinary Least Squares regression to determine which independent variables (sample type, bacterial count, read count) explain the dependent outcome variable (the immune score). A p-value ≤ 0.05 was considered statistically significant.

**Table S5 – Bacterial counts impacting specific immune cells across the seven methods that score immune cell infiltration, excluding paired normal samples from the control group.** NA = Not applicable because the method does not report that type of immune cell. ns = not significant. * = p<0.05, ** = p<0.01, *** p<0.001, **** p<0.0001. Statistical comparison was carried out with Ordinary Least Squares regression to determine which independent variables (sample type, bacterial count, read count) explain the dependent outcome variable (the immune score). Highlighted rows indicate immune cell sub-populations only affected in CRC tissue by bacterial counts. A p-value ≤ 0.05 was considered statistically significant.

**Table S6 – *Bacteroides fragilis* impacting specific immune cells across the seven methods that score immune cell infiltration excluding paired normal samples from the control group.** NA = Not applicable because the method does not report that type of immune cell. ns = not significant. * = p<0.05, ** = p<0.01, *** p<0.001. Statistical comparison was carried out with Ordinary Least Squares regression to determine which independent variables (sample type, bacterial count, read count) explain the dependent outcome variable (the immune score). Highlighted rows indicate immune cell sub-populations only affected in CRC tissue by bacterial counts. A p-value ≤ 0.05 was considered statistically significant.

**Table S7 – *Fusobacterium nucleatum* impacting specific immune cells across the seven methods that score immune cell infiltration excluding paired normal samples from the control group.** NA = Not applicable because the method does not report that type of immune cell. ns = not significant. * = p<0.05. Statistical comparison was carried out with Ordinary Least Squares regression to determine which independent variables (sample type, bacterial count, read count) explain the dependent outcome variable (the immune score). A p-value ≤ 0.05 was considered statistically significant.

**Figure S1 – Noise filtering and distribution of counts assigned to Eukaryota, bacteria, archaea, viruses, *Fusobacterium nucleatum,* and *Bacteroides fragilis.* A)** Histogram showing the counts distribution over log-10 transformed scaled counts. The red arrow indicates the intersection between the populations, and all scaled counts < log-0.9 (indicated by red arrow) were set to 0 to remove noise. **B+C+D+E+F+G)** Normal distribution of scaled kingdom counts (y-axis) presented per sample across groups for Eukaryota (A), Bacteria (B), Archaea (C), virus (D), *F. nucleatum* (F), and *B. fragilis* (G).

**Figure S2 – bacterial biomass was not associated with tumour staging, lymph node metastasis, or distant metastasis. A)** Logarithmic (log) transformed bacterial biomass measured in cubic micrometers (µm^3^) according to tumour stage (T1-4). **B+C+D)** Log-transformed bacterial biomass measured in µm^3^ according to distant (B) metastasis (M), Lymph node (C) metastasis (N), or both (D). All biomass measurements were measured with the Imaris software through thresholding of fluorescence intensity. **E+F+G)** Prevalence of *Fusobacterium spp.* compared with the number of patients with distant (E) metastasis (M), Lymph node (F) metastasis (N), or both (G**). H+I)** Prevalence of *Bacteroides fragilis* compared with the number of patients with lymph node (H) metastasis (N) or distant (I) metastasis (M**). J+K)** Number of patients and tumour staging (T1-4) compared to the prevalence of *B. fragilis* (J) and *Fusobacterium spp.* (K). Statistical comparison was carried out with one-way ANOVA **(A),** unpaired t-test **(B+C)**, Fisher’s exact t-test **(A+B+C+D),** and chi-square test **(E+F)**. Bars represent standard deviation (SD); a p-value ≤ 0.05 was considered statistically significant.

**Figure S3 – Principal-component analysis of normalized expression data. A+B)** The clustering of CRC, healthy, and paired normal tissue samples according to sequencing depth (deep vs. shallow) is presented as a heatmap (A) and 2D scatterplot (B). **C)** All data is colored according to groups. **D)** All data is colored according to sequencing depth. **E)** CRC samples are colored by sequencing depth. **F)** CRC samples are colored according to the presence of *F. nucleatum*. Samples with high *F. nucleatum* counts were defined as those samples departing from the normal distribution in Figure S1, whereas low were those that followed the normal distribution. **G)** CRC samples are colored according to the presence of *B. fragilis*. Samples with high *B. fragilis* counts were defined as those departing from the normal distribution in Figure S1, whereas low were those following the normal distribution.

**Figure S4 – Differentially expressed genes and enriched biological pathways in CRC and non-CRC. A)** MA plot showing the distribution of significantly differentiated genes between CRC and Non-CRC (healthy and paired samples). Coloring highlights the 20 most significant DEGs with an adjusted p-value less than 0.05 and absolute log2 fold-change >2. Coloring highlights genes with an adjusted p-value less than 0.05 and absolute log2 fold-change >2. **B)** Pathways demonstrating significant enrichment of differentially expressed genes (Fishers exact test) for CRC or non-CRC. The Kyoto Encyclopedia of Genes and Genomes (KEGG), Pathway Interaction Database (PID), and REACTOME (a database of reactions, pathways, and biological processes) databases were used to identify pathways.

**Figure S5 – Clustering of samples according to immune cell infiltration 2. A+B)** Heatmaps showing immune cell profiles in CRC and healthy tissue, presented as normalized scores from the concensus_tme, xcell, mcp_counter, and timer immune scoring systems (A), and fractions from the quantisec, epic, and estimate immune scoring systems (B). Coloring from yellow (−4) to purple (4) indicates the degree of infiltration, where purple is high infiltration.

## REFERENCES

1. Zhao L, Grimes SM, Greer SU, et al. Characterization of the consensus mucosal microbiome of colorectal cancer. NAR Cancer 2021;3(4) doi: 10.1093/narcan/zcab049

2. Yu J, Feng Q, Wong SH, et al. Metagenomic analysis of faecal microbiome as a tool towards targeted non-invasive biomarkers for colorectal cancer. Gut 2017;66(1):70–78. doi: 10.1136/gutjnl-2015-309800 [published Online First: 2015/09/27]

3. Yachida S, Mizutani S, Shiroma H, et al. Metagenomic and metabolomic analyses reveal distinct stage-specific phenotypes of the gut microbiota in colorectal cancer. Nature medicine 2019;25(6):968–76. doi: 10.1038/s41591-019-0458-7 [published Online First: 2019/06/07]

4. Amitay EL, Krilaviciute A, Brenner H. Systematic review: Gut microbiota in fecal samples and detection of colorectal neoplasms. Gut Microbes 2018;9(4):293–307. doi: 10.1080/19490976.2018.1445957 [published Online First: 2018/03/16]

5. Janney A, Powrie F, Mann EH. Host-microbiota maladaptation in colorectal cancer. Nature 2020;585(7826):509-17. doi: 10.1038/s41586-020-2729-3 [published Online First: 2020/09/25]

6. Coussens LM, Werb Z. Inflammation and cancer. Nature 2002;420(6917):860-7. doi: 10.1038/nature01322 [published Online First: 2002/12/20]

7. Tjalsma H, Boleij A, Marchesi JR, et al. A bacterial driver–passenger model for colorectal cancer: beyond the usual suspects. Nature Reviews Microbiology 2012;10(8):575–82. doi: 10.1038/nrmicro2819

8. Bennedsen ALB, Furbo S, Bjarnsholt T, et al. The gut microbiota can orchestrate the signaling pathways in colorectal cancer. APMIS: acta pathologica, microbiologica, et immunologica Scandinavica 2022;130(3):121–39. doi: 10.1111/apm.13206 [published Online First: 2022/01/11]

9. Geis AL, Fan H, Wu X, et al. Regulatory T-cell Response to Enterotoxigenic Bacteroides fragilis Colonization Triggers IL17-Dependent Colon Carcinogenesis. Cancer Discov 2015;5(10):1098–109. doi: 10.1158/2159-8290.Cd-15-0447 [published Online First: 2015/07/24]

10. Wu J, Li Q, Fu X. Fusobacterium nucleatum Contributes to the Carcinogenesis of Colorectal Cancer by Inducing Inflammation and Suppressing Host Immunity. Translational oncology 2019;12(6):846–51. doi: 10.1016/j.tranon.2019.03.003 [published Online First: 2019/04/16]

11. Kostic AD, Chun E, Robertson L, et al. Fusobacterium nucleatum potentiates intestinal tumorigenesis and modulates the tumor-immune microenvironment. Cell Host Microbe 2013;14(2):207–15. doi: 10.1016/j.chom.2013.07.007 [published Online First: 2013/08/21]

12. Gur C, Maalouf N, Shhadeh A, et al. Fusobacterium nucleatum supresses anti-tumor immunity by activating CEACAM1. Oncoimmunology 2019;8(6):e1581531. doi: 10.1080/2162402x.2019.1581531 [published Online First: 2019/05/10]

13. Xue Y, Xiao H, Guo S, et al. Indoleamine 2,3-dioxygenase expression regulates the survival and proliferation of Fusobacterium nucleatum in THP-1-derived macrophages. Cell Death Dis 2018;9(3):355. doi: 10.1038/s41419-018-0389-0 [published Online First: 2018/03/04]

14. Brennan CA, Garrett WS. Gut Microbiota, Inflammation, and Colorectal Cancer. Annu Rev Microbiol 2016;70:395-411. doi: 10.1146/annurev-micro-102215-095513 [published Online First: 2016/09/09]

15. Fritz BG, Kirkegaard JB, Nielsen CH, et al. Transcriptomic fingerprint of bacterial infection in lower extremity ulcers. APMIS: acta pathologica, microbiologica, et immunologica Scandinavica 2022;130(8):524–34. doi: 10.1111/apm.13234 [published Online First: 2022/05/15]

16. Cornforth DM, Dees JL, Ibberson CB, et al. Pseudomonas aeruginosa transcriptome during human infection. Proc Natl Acad Sci U S A 2018;115(22):E5125–e34. doi: 10.1073/pnas.1717525115 [published Online First: 2018/05/16]

17. Saus E, Iraola-Guzmán S, Willis JR, et al. Microbiome and colorectal cancer: Roles in carcinogenesis and clinical potential. Molecular aspects of medicine 2019;69:93–106. doi: 10.1016/j.mam.2019.05.001 [published Online First: 2019/05/15]

18. Aitmanaitė L, Širmonaitis K, Russo G. Microbiomes, Their Function, and Cancer: How Metatranscriptomics Can Close the Knowledge Gap. International journal of molecular sciences 2023;24(18) doi: 10.3390/ijms241813786 [published Online First: 2023/09/28]

19. Rigottier-Gois L, Rochet V, Garrec N, et al. Enumeration of Bacteroides species in human faeces by fluorescent in situ hybridisation combined with flow cytometry using 16S rRNA probes. Systematic and applied microbiology 2003;26(1):110–8. doi: 10.1078/072320203322337399 [published Online First: 2003/05/16]

20. Valm AM, Welch JLM, Rieken CW, et al. Systems-level analysis of microbial community organization through combinatorial labeling and spectral imaging. Proceedings of the National Academy of Sciences 2011;108(10):4152–57. doi: 10.1073/pnas.1101134108

21. Bay L, Kragh KN, Eickhardt SR, et al. Bacterial Aggregates Establish at the Edges of Acute Epidermal Wounds. Adv Wound Care (New Rochelle) 2018;7(4):105–13. doi: 10.1089/wound.2017.0770 [published Online First: 2018/04/21]

22. Stender H, Mollerup TA, Lund K, et al. Direct detection and identification of Mycobacterium tuberculosis in smear-positive sputum samples by fluorescence in situ hybridization (FISH) using peptide nucleic acid (PNA) probes. The international journal of tuberculosis and lung disease: the official journal of the International Union against Tuberculosis and Lung Disease 1999;3(9):830–7. [published Online First: 1999/09/17]

23. Kragh KN, Alhede M, Kvich L, et al. Into the well-A close look at the complex structures of a microtiter biofilm and the crystal violet assay. Biofilm 2019;1:100006. doi: 10.1016/j.bioflm.2019.100006 [published Online First: 2019/09/12]

24. Klopfleisch R. Multiparametric and semiquantitative scoring systems for the evaluation of mouse model histopathology--a systematic review. BMC Vet Res 2013;9:123. doi: 10.1186/1746-6148-9-123 [published Online First: 2013/06/27]

25. Kolpen M, Kragh KN, Enciso JB, et al. Bacterial biofilms predominate in both acute and chronic human lung infections. Thorax 2022;77(10):1015–22. doi: 10.1136/thoraxjnl-2021-217576 [published Online First: 2022/01/13]

26. Martin M. Cutadapt removes adapter sequences from high-throughput sequencing reads. 2011 2011;17(1):3. doi: 10.14806/ej.17.1.200 [published Online First: 2011-08-02]

27. Kopylova E, Noé L, Touzet H. SortMeRNA: fast and accurate filtering of ribosomal RNAs in metatranscriptomic data. Bioinformatics 2012;28(24):3211–7. doi: 10.1093/bioinformatics/bts611 [published Online First: 2012/10/17]

28. Li H. Aligning sequence reads, clone sequences and assembly contigs with BWA-MEM. arXiv: Genomics 2013

29. Liao Y, Smyth GK, Shi W. The Subread aligner: fast, accurate and scalable read mapping by seed-and-vote. Nucleic Acids Res 2013;41(10):e108. doi: 10.1093/nar/gkt214 [published Online First: 2013/04/06]

30. Wood DE, Lu J, Langmead B. Improved metagenomic analysis with Kraken 2. Genome Biology 2019;20(1):257. doi: 10.1186/s13059-019-1891-0

31. Lu J, Breitwieser F.P., Thielen P., et al. Bracken: estimating species abundance in metagenomics data. PeerJ Computer Science 2017;3:e104 doi: 10.7717/peerj-cs.104

32. Durinck S, Moreau Y, Kasprzyk A, et al. BioMart and Bioconductor: a powerful link between biological databases and microarray data analysis. Bioinformatics 2005;21(16):3439–40. doi: 10.1093/bioinformatics/bti525 [published Online First: 2005/08/06]

33. Priya S, Burns MB, Ward T, et al. Identification of shared and disease-specific host gene–microbiome associations across human diseases using multi-omic integration. Nature Microbiology 2022;7(6):780–95. doi: 10.1038/s41564-022-01121-z

34. Nederlof I, De Bortoli D, Bareche Y, et al. Comprehensive evaluation of methods to assess overall and cell-specific immune infiltrates in breast cancer. Breast Cancer Res 2019;21(1):151. doi: 10.1186/s13058-019-1239-4 [published Online First: 2019/12/28]

35. Finotello F, Trajanoski Z. Quantifying tumor-infiltrating immune cells from transcriptomics data. Cancer immunology, immunotherapy: CII 2018;67(7):1031–40. doi: 10.1007/s00262-018-2150-z<otherinfo> [published Online First: 2018/03/16]</otherinfo>

36. Sturm G, Finotello F, List M. Immunedeconv: An R Package for Unified Access to Computational Methods for Estimating Immune Cell Fractions from Bulk RNA-Sequencing Data. *Methods in molecular biology (Clifton*, NJ) 2020;2120:223–32. doi: 10.1007/978-1-0716-0327-7_16 [published Online First: 2020/03/04]

37. Ritchie ME, Phipson B, Wu D, et al. limma powers differential expression analyses for RNA-sequencing and microarray studies. Nucleic Acids Research 2015;43(7):e47–e47. doi: 10.1093/nar/gkv007

38. Dejea CM, Wick EC, Hechenbleikner EM, et al. Microbiota organization is a distinct feature of proximal colorectal cancers. Proc Natl Acad Sci U S A 2014;111(51):18321–6. doi: 10.1073/pnas.1406199111 [published Online First: 2014/12/10]

39. Lima BP, Shi W, Lux R. Identification and characterization of a novel Fusobacterium nucleatum adhesin involved in physical interaction and biofilm formation with Streptococcus gordonii. MicrobiologyOpen 2017;6(3):e00444. doi: 10.1002/mbo3.444

40. Coppenhagen-Glazer S, Sol A, Abed J, et al. Fap2 of Fusobacterium nucleatum is a galactose-inhibitable adhesin involved in coaggregation, cell adhesion, and preterm birth. Infection and immunity 2015;83(3):1104–13. doi: 10.1128/iai.02838-14 [published Online First: 2015/01/07]

41. Kaplan CW, Lux R, Haake SK, et al. The Fusobacterium nucleatum outer membrane protein RadD is an arginine-inhibitable adhesin required for inter-species adherence and the structured architecture of multispecies biofilm. Mol Microbiol 2009;71(1):35–47. doi: 10.1111/j.1365-2958.2008.06503.x [published Online First: 2008/11/15]

42. Liu PF, Shi W, Zhu W, et al. Vaccination targeting surface FomA of Fusobacterium nucleatum against bacterial co-aggregation: Implication for treatment of periodontal infection and halitosis. Vaccine 2010;28(19):3496–505. doi: 10.1016/j.vaccine.2010.02.047 [published Online First: 2010/03/02]

43. Kinross J, Mirnezami R, Alexander J, et al. A prospective analysis of mucosal microbiome-metabonome interactions in colorectal cancer using a combined MAS 1HNMR and metataxonomic strategy. Scientific Reports 2017;7(1):8979. doi: 10.1038/s41598-017-08150-3

44. Drewes JL, White JR, Dejea CM, et al. High-resolution bacterial 16S rRNA gene profile meta-analysis and biofilm status reveal common colorectal cancer consortia. NPJ biofilms and microbiomes 2017;3:34. doi: 10.1038/s41522-017-0040-3 [published Online First: 2017/12/08]

45. Bullman S, Pedamallu CS, Sicinska E, et al. Analysis of Fusobacterium persistence and antibiotic response in colorectal cancer. Science 2017;358(6369):1443–48. doi: 10.1126/science.aal5240 [published Online First: 2017/11/25]

46. Zhao L, Grimes SM, Greer SU, et al. Characterization of the consensus mucosal microbiome of colorectal cancer. NAR Cancer 2021;3(4):zcab049. doi: 10.1093/narcan/zcab049 [published Online First: 2022/01/07]

47. Nakatsu G, Li X, Zhou H, et al. Gut mucosal microbiome across stages of colorectal carcinogenesis. Nature Communications 2015;6(1):8727. doi: 10.1038/ncomms9727

48. Purcell RV, Visnovska M, Biggs PJ, et al. Distinct gut microbiome patterns associate with consensus molecular subtypes of colorectal cancer. Sci Rep 2017;7(1):11590. doi: 10.1038/s41598-017-11237-6 [published Online First: 2017/09/16]

49. Tahara T, Yamamoto E, Suzuki H, et al. Fusobacterium in colonic flora and molecular features of colorectal carcinoma. Cancer Res 2014;74(5):1311–8. doi: 10.1158/0008-5472.can-13-1865 [published Online First: 2014/01/05]

50. Chen Y, Huang Z, Tang Z, et al. More Than Just a Periodontal Pathogen-the Research Progress on Fusobacterium nucleatum. Front Cell Infect Microbiol 2022;12:815318. doi: 10.3389/fcimb.2022.815318 [published Online First: 2022/02/22]

51. Saffarian A, Mulet C, Regnault B, et al. Crypt- and Mucosa-Associated Core Microbiotas in Humans and Their Alteration in Colon Cancer Patients. mBio 2019;10(4) doi: 10.1128/mBio.01315-19 [published Online First: 2019/07/18]

52. Clay SL, Fonseca-Pereira D, Garrett WS. Colorectal cancer: the facts in the case of the microbiota. The Journal of clinical investigation 2022;132(4) doi: 10.1172/jci155101 [published Online First: 2022/02/16]

53. Boleij A, Hechenbleikner EM, Goodwin AC, et al. The Bacteroides fragilis toxin gene is prevalent in the colon mucosa of colorectal cancer patients. Clin Infect Dis 2015;60(2):208–15. doi: 10.1093/cid/ciu787 [published Online First: 2014/10/12]

54. Brennan CA, Clay SL, Lavoie SL, et al. Fusobacterium nucleatum drives a pro-inflammatory intestinal microenvironment through metabolite receptor-dependent modulation of IL-17 expression. Gut Microbes 2021;13(1):1987780. doi: 10.1080/19490976.2021.1987780 [published Online First: 2021/11/17]

