## Supplemental tables 1-7 and figures 1-5 for "Biofilms and core pathogens shape the tumour microenvironment and immune phenotype in colorectal cancer"

**Supplemental information**

| Characteristics | CRC, N = 37 (%) | Healthy, N = 40 (%) | P-value |
|---|---|---|---|
| Sex, male | 22 (61.1) | 25 (62.5) | >0.99 |
| Age, median (range)<br>Weight (Kg.), median (range)<br>BMI, median (range)<br>Missing information (BMI) | 76 (47 – 90)<br>74 (46 – 119)<br>24.8 (16.7 – 41.2)<br>2 | 68.5 (46-86)<br>76 (49-121)<br>25.8 (17.6-42.9)<br>2 | 0.05<br>0.71<br>0.40 |
| Anatomic location of sampling<br>Left-sided<br>Right-sided<br>Missing information | <br>28 (75.7)<br>9 (24.3) | <br>22 (56.4)<br>17 (43.6)<br>1 | <br>0.09 |
| T stage<br>1<br>2<br>3<br>4<br>Missing information | <br>4 (10.8)<br>7 (18.9)<br>13 (35.1)<br>5 (13.5)<br>8 | <br>N/A<br>N/A<br>N/A<br>N/A | <br>N/A<br>N/A<br>N/A<br>N/A |
| Lymph node metastases (N)<br>Distant metastases (M)<br>Missing information (N/M)<br>pMMR/dMMR | 12 (32.4)<br>7 (18.9)<br>3/2<br>35/2 | N/A<br>N/A<br><br>N/A | N/A<br>N/A<br><br>N/A |
| ASA score<br>1<br>2<br>3<br>Missing information | <br>2 (5.4)<br>24 (64.9)<br>11 (29.7) | <br>6 (15.0)<br>20 (50.0)<br>1 (2.5)<br>13 | <br><0.01 |
| Smoking<br>Yes<br>No<br>Previously<br>Missing information | <br>9 (24.3)<br>12 (32.4)<br>15 (40.5)<br>1 | <br>3 (7.5)<br>14 (35)<br>15 (37)<br>8 | <br>0.46* |

**Table S1 - Characteristics of included patients with CRC and healthy persons**

ASA, American Society of Anesthesiologists. BMI, Body Mass Index (Kg/m^2). DM, Diabetes mellitus. *Current and previous smoking has been pooled for statistical analysis. Continuous data was tested with a two-sided student t-test or Mann-Whitney Test, and categorical data was tested with a chi-square test. A p-value ≤ 0.05 was considered statistically significant.

| Cell_type | Estimate frac | Estimate score | timer | consensus_tme | xcell | mcp_counter | epic | quantiseq |
|---|---|---|---|---|---|---|---|---|
| B cell | NA | NA | ns | ns | ns | ns | ns | NA |
| B cell memory | NA | NA | NA | NA | ns | NA | NA | NA |
| B cell naive | NA | NA | NA | NA | ns | NA | NA | NA |
| B cell plasma | NA | NA | NA | NA | ns | NA | NA | NA |
| Cancer associated fibroblast | NA | NA | NA | ns | ns | ns | ns | NA |
| Class-switched memory B cell | NA | NA | NA | NA | ns | NA | NA | NA |
| Common lymphoid progenitor | NA | NA | NA | NA | ns | NA | NA | NA |
| Common myeloid progenitor | NA | NA | NA | NA | ns | NA | NA | NA |
| cytotoxicity score | NA | NA | NA | ns | NA | ns | NA | NA |
| Endothelial cell | NA | NA | NA | ** | **** | *** | * | NA |
| Eosinophil | NA | NA | NA | ns | ns | NA | NA | NA |
| Granulocyte-monocyte progenitor | NA | NA | NA | NA | ns | NA | NA | NA |
| Hematopoietic stem cell | NA | NA | NA | NA | *** | NA | NA | NA |
| immune score | NA | ns | NA | ns | ns | NA | NA | NA |
| Macrophage | NA | NA | ns | ns | ns | NA | ns | NA |
| Macrophage M1 | NA | NA | NA | ns | ns | NA | NA | NA |
| Macrophage M2 | NA | NA | NA | ns | ns | NA | NA | NA |
| **Macrophage/Monocyte** | **NA** | **NA** | **NA** | **NA** | **NA** | ** | **NA** | **NA** |
| Mast cell | NA | NA | NA | NA | ns | NA | NA | NA |
| MAST cell | NA | NA | NA | ns | NA | NA | NA | NA |
| microenvironment score | NA | NA | NA | NA | ns | NA | NA | NA |
| Monocyte | NA | NA | NA | ns | * | ** | NA | NA |
| Myeloid dendritic cell | NA | NA | ** | ns | ns | ns | NA | NA |
| **Myeloid dendritic cell activated** | **NA** | **NA** | **NA** | **NA** | * | **NA** | **NA** | **NA** |
| Neutrophil | NA | NA | ** | ns | * | ns | NA | NA |
| NK cell | NA | NA | NA | ns | ns | ns | ns | NA |
| Plasmacytoid dendritic cell | NA | NA | NA | NA | ns | NA | NA | NA |
| stroma score | NA | ** | NA | NA | * | NA | NA | NA |
| T cell | NA | NA | NA | NA | NA | ns | NA | NA |
| T cell CD4+ | NA | NA | ns | ns | NA | NA | ns | NA |
| T cell CD4+ (non-regulatory) | NA | NA | NA | NA | ns | NA | NA | NA |
| T cell CD4+ central memory | NA | NA | NA | NA | ns | NA | NA | NA |
| T cell CD4+ effector memory | NA | NA | NA | NA | ns | NA | NA | NA |
| T cell CD4+ memory | NA | NA | NA | NA | ns | NA | NA | NA |

| | | | | | | | | |
|---|---|---|---|---|---|---|---|---|
| T cell CD4+ naive | NA | NA | NA | NA | ns | NA | NA | NA |
| T cell CD4+ Th1 | NA | NA | NA | NA | ns | NA | NA | NA |
| T cell CD4+ Th2 | NA | NA | NA | NA | ns | NA | NA | NA |
| T cell CD8+ | NA | NA | ns | ns | ns | ns | * | NA |
| T cell CD8+ central memory | NA | NA | NA | NA | ns | NA | NA | NA |
| T cell CD8+ effector memory | NA | NA | NA | NA | ns | NA | NA | NA |
| T cell CD8+ naive | NA | NA | NA | NA | ns | NA | NA | NA |
| T cell gamma delta | NA | NA | NA | ns | ns | NA | NA | NA |
| T cell NK | NA | NA | NA | NA | ns | NA | NA | NA |
| **T cell regulatory (Tregs)** | **NA** | **NA** | **NA** | **ns** | ******* | **NA** | **NA** | **NA** |
| **tumor purity fraction** | ******* | **NA** | **NA** | **NA** | **NA** | **NA** | **NA** | **NA** |
| **tumor purity score** | **NA** | ***** | **NA** | **NA** | **NA** | **NA** | **NA** | **NA** |
| uncharacterized cell | NA | NA | NA | NA | NA | NA | ns | NA |

**Table S2 – Bacterial counts impacting specific immune cells across the seven methods that score immune cell infiltration.**

NA = Not applicable because the method does not report that type of immune cell. ns = not significant. * = $p<0.05$, ** = $p<0.01$, *** $p<0.001$, **** $p<0.0001$. Statistical comparison was carried out with Ordinary Least Squares regression to determine which independent variables (sample type, bacterial count, read count) explain the dependent outcome variable (the immune score). Highlighted rows with bold text indicate immune cell sub-populations only affected in CRC tissue by bacterial counts. A p-value $\leq 0.05$ was considered statistically significant.

| Cell_type | Estimate_frac | Estimate_score | Timer | consensus_tme | xcell | mcp_counter | epic | quantiseq |
|---|---|---|---|---|---|---|---|---|
| B cell | NA | NA | ns | ns | ns | ns | ns | NA |
| B cell memory | NA | NA | NA | NA | ns | NA | NA | NA |
| B cell naive | NA | NA | NA | NA | ns | NA | NA | NA |
| B cell plasma | NA | NA | NA | NA | ns | NA | NA | NA |
| Cancer associated fibroblast | NA | NA | NA | ns | ns | ns | ns | NA |
| Class-switched memory B cell | NA | NA | NA | NA | ns | NA | NA | NA |
| Common lymphoid progenitor | NA | NA | NA | NA | ns | NA | NA | NA |
| Common myeloid progenitor | NA | NA | NA | NA | ns | NA | NA | NA |
| cytotoxicity score | NA | NA | NA | ns | NA | ns | NA | NA |
| Endothelial cell | NA | NA | NA | ns | ns | ns | ns | NA |
| Eosinophil | NA | NA | NA | ns | ns | NA | NA | NA |
| Granulocyte-monocyte progenitor | NA | NA | NA | NA | ns | NA | NA | NA |
| Hematopoietic stem cell | NA | NA | NA | NA | ns | NA | NA | NA |
| immune score | NA | ns | NA | ns | ns | NA | NA | NA |
| Macrophage | NA | NA | ns | ns | ns | NA | ns | NA |
| Macrophage M1 | NA | NA | NA | ns | ns | NA | NA | NA |
| Macrophage M2 | NA | NA | NA | ns | ns | NA | NA | NA |
| Macrophage/Monocyte | NA | NA | NA | NA | NA | ns | NA | NA |
| Mast cell | NA | NA | NA | NA | ns | NA | NA | NA |
| MAST cell | NA | NA | NA | ns | NA | NA | NA | NA |
| microenvironment score | NA | NA | NA | NA | ns | NA | NA | NA |
| Monocyte | NA | NA | NA | ns | ns | ns | NA | NA |
| Myeloid dendritic cell | NA | NA | ns | ns | ns | ns | NA | NA |
| Myeloid dendritic cell activated | NA | NA | NA | NA | ns | NA | NA | NA |
| **Neutrophil** | **NA** | **NA** | **ns** | **ns** | ***** | **ns** | **NA** | **NA** |
| NK cell | NA | NA | NA | ns | ns | ns | ns | NA |
| Plasmacytoid dendritic cell | NA | NA | NA | NA | ns | NA | NA | NA |
| stroma score | NA | ns | NA | NA | ns | NA | NA | NA |
| T cell | NA | NA | NA | NA | NA | ns | NA | NA |
| T cell CD4+ | NA | NA | ns | ns | NA | NA | ns | NA |
| T cell CD4+ (non-regulatory) | NA | NA | NA | NA | ns | NA | NA | NA |
| T cell CD4+ central memory | NA | NA | NA | NA | ns | NA | NA | NA |
| T cell CD4+ effector memory | NA | NA | NA | NA | ns | NA | NA | NA |
| T cell CD4+ memory | NA | NA | NA | NA | ns | NA | NA | NA |

| | | | | | | | | |
|---|---|---|---|---|---|---|---|---|
| T cell CD4+ naive | NA | NA | NA | NA | ns | NA | NA | NA |
| T cell CD4+ Th1 | NA | NA | NA | NA | ns | NA | NA | NA |
| T cell CD4+ Th2 | NA | NA | NA | NA | ns | NA | NA | NA |
| T cell CD8+ | NA | NA | ns | ns | ns | ns | ns | NA |
| T cell CD8+ central memory | NA | NA | NA | NA | ns | NA | NA | NA |
| T cell CD8+ effector memory | NA | NA | NA | NA | ns | NA | NA | NA |
| T cell CD8+ naive | NA | NA | NA | NA | ns | NA | NA | NA |
| T cell gamma delta | NA | NA | NA | ns | ns | NA | NA | NA |
| T cell NK | NA | NA | NA | NA | ns | NA | NA | NA |
| T cell regulatory (Tregs) | NA | NA | NA | ns | ns | NA | NA | NA |
| **tumor purity fraction** | ***** | **NA** | **NA** | **NA** | **NA** | **NA** | **NA** | **NA** |
| tumor purity score | NA | ns | NA | NA | NA | NA | NA | NA |
| uncharacterized cell | NA | NA | NA | NA | NA | NA | ns | NA |

**Table S3 – *Bacteroides fragilis* impacting specific immune cells across the seven methods that score immune cell infiltration.**

NA = Not applicable because the method does not report that type of immune cell. ns = not significant. * = $p<0.05$. Statistical comparison was carried out with Ordinary Least Squares regression to determine which independent variables (sample type, bacterial count, read count) explain the dependent outcome variable (the immune score). Highlighted rows with bold text indicate immune cell sub-populations only affected in CRC tissue by *B. fragilis*. A p-value $\leq 0.05$ was considered statistically significant.

| Cell_type | Estimate_frac | Estimate_score | Timer | consensus_tme | xcell | mcp_counter | epic | quantiseq |
|---|---|---|---|---|---|---|---|---|
| B cell | NA | NA | ns | ns | ns | ns | ns | NA |
| B cell memory | NA | NA | NA | NA | ns | NA | NA | NA |
| B cell naive | NA | NA | NA | NA | ns | NA | NA | NA |
| B cell plasma | NA | NA | NA | NA | ns | NA | NA | NA |
| Cancer associated fibroblast | NA | NA | NA | ns | ns | ns | ns | NA |
| Class-switched memory B cell | NA | NA | NA | NA | ns | NA | NA | NA |
| Common lymphoid progenitor | NA | NA | NA | NA | ns | NA | NA | NA |
| Common myeloid progenitor | NA | NA | NA | NA | ns | NA | NA | NA |
| cytotoxicity score | NA | NA | NA | ns | NA | ns | NA | NA |
| Endothelial cell | NA | NA | NA | ns | ns | ns | ns | NA |
| Eosinophil | NA | NA | NA | ns | ns | NA | NA | NA |
| Granulocyte-monocyte progenitor | NA | NA | NA | NA | ns | NA | NA | NA |
| Hematopoietic stem cell | NA | NA | NA | NA | ns | NA | NA | NA |
| immune score | NA | ns | NA | ns | ns | NA | NA | NA |
| Macrophage | NA | NA | ns | ns | ns | NA | ns | NA |
| Macrophage M1 | NA | NA | NA | ns | ns | NA | NA | NA |
| Macrophage M2 | NA | NA | NA | ns | ns | NA | NA | NA |
| Macrophage/Monocyte | NA | NA | NA | NA | NA | ns | NA | NA |
| Mast cell | NA | NA | NA | NA | ns | NA | NA | NA |
| MAST cell | NA | NA | NA | ns | NA | NA | NA | NA |
| microenvironment score | NA | NA | NA | NA | ns | NA | NA | NA |
| Monocyte | NA | NA | NA | ns | ns | ns | NA | NA |
| Myeloid dendritic cell | NA | NA | ns | ns | ns | ns | NA | NA |
| Myeloid dendritic cell activated | NA | NA | NA | NA | ns | NA | NA | NA |
| Neutrophil | NA | NA | ns | ns | ns | ns | NA | NA |
| NK cell | NA | NA | NA | ns | ns | ns | ns | NA |
| Plasmacytoid dendritic cell | NA | NA | NA | NA | ns | NA | NA | NA |
| stroma score | NA | ns | NA | NA | ns | NA | NA | NA |
| T cell | NA | NA | NA | NA | NA | ns | NA | NA |
| T cell CD4+ | NA | NA | ns | ns | NA | NA | ns | NA |
| T cell CD4+ (non-regulatory) | NA | NA | NA | NA | ns | NA | NA | NA |
| T cell CD4+ central memory | NA | NA | NA | NA | ns | NA | NA | NA |
| T cell CD4+ effector memory | NA | NA | NA | NA | ns | NA | NA | NA |
| T cell CD4+ memory | NA | NA | NA | NA | ns | NA | NA | NA |

| | | | | | | | | |
|---|---|---|---|---|---|---|---|---|
| T cell CD4+ naive | NA | NA | NA | NA | ns | NA | NA | NA |
| T cell CD4+ Th1 | NA | NA | NA | NA | ns | NA | NA | NA |
| T cell CD4+ Th2 | NA | NA | NA | NA | ns | NA | NA | NA |
| T cell CD8+ | NA | NA | ns | ns | ns | ns | ns | NA |
| T cell CD8+ central memory | NA | NA | NA | NA | ns | NA | NA | NA |
| T cell CD8+ effector memory | NA | NA | NA | NA | ns | NA | NA | NA |
| T cell CD8+ naive | NA | NA | NA | NA | ns | NA | NA | NA |
| T cell gamma delta | NA | NA | NA | ns | ns | NA | NA | NA |
| T cell NK | NA | NA | NA | NA | ns | NA | NA | NA |
| T cell regulatory (Tregs) | NA | NA | NA | ns | ns | NA | NA | NA |
| tumor purity fraction | ns | NA | NA | NA | NA | NA | NA | NA |
| tumor purity score | NA | ns | NA | NA | NA | NA | NA | NA |
| uncharacterized cell | NA | NA | NA | NA | NA | NA | ns | NA |

**Table S4 – *Fusobacterium nucleatum* impacting specific immune cells across the seven methods that score immune cell infiltration.**

NA = Not applicable because the method does not report that type of immune cell. ns = not significant. Statistical comparison was carried out with Ordinary Least Squares regression to determine which independent variables (sample type, bacterial count, read count) explain the dependent outcome variable (the immune score). A p-value $\leq 0.05$ was considered statistically significant.

| Cell_type | Estimate _frac | Estimate _score | Timer | Consensus _tme | xcell | Mcp _counter | epic | quantiseq |
|---|---|---|---|---|---|---|---|---|
| B cell | NA | NA | ns | ns | ns | ns | ns | NA |
| B cell memory | NA | NA | NA | NA | ns | NA | NA | NA |
| B cell naive | NA | NA | NA | NA | ns | NA | NA | NA |
| B cell plasma | NA | NA | NA | NA | ns | NA | NA | NA |
| Cancer associated fibroblast | NA | NA | NA | ns | ns | ns | ns | NA |
| Class-switched memory B cell | NA | NA | NA | NA | ns | NA | NA | NA |
| Common lymphoid progenitor | NA | NA | NA | NA | ns | NA | NA | NA |
| Common myeloid progenitor | NA | NA | NA | NA | ns | NA | NA | NA |
| cytotoxicity score | NA | NA | NA | ns | NA | ns | NA | NA |
| Endothelial cell | NA | NA | NA | ** | *** | ** | * | NA |
| **Eosinophil** | **NA** | **NA** | **NA** | ***** | **ns** | **NA** | **NA** | **NA** |
| Granulocyte-monocyte progenitor | NA | NA | NA | NA | ns | NA | NA | NA |
| **Hematopoietic stem cell** | **NA** | **NA** | **NA** | **NA** | ****** | **NA** | **NA** | **NA** |
| **immune score** | **NA** | ****** | **NA** | ***** | **ns** | **NA** | **NA** | **NA** |
| Macrophage | NA | NA | ns | * | ns | NA | ns | NA |
| Macrophage M1 | NA | NA | NA | ns | ns | NA | NA | NA |
| **Macrophage M2** | **NA** | **NA** | **NA** | ***** | ***** | **NA** | **NA** | **NA** |
| **Macrophage/Monocyte** | **NA** | **NA** | **NA** | **NA** | **NA** | ****** | **NA** | **NA** |
| Mast cell | NA | NA | NA | NA | ns | NA | NA | NA |
| MAST cell | NA | NA | NA | ns | NA | NA | NA | NA |
| **microenvironment score** | **NA** | **NA** | **NA** | **NA** | ***** | **NA** | **NA** | **NA** |
| **Monocyte** | **NA** | **NA** | **NA** | ***** | **ns** | ****** | **NA** | **NA** |
| Myeloid dendritic cell | NA | NA | *** | * | ns | ns | NA | NA |
| **Myeloid dendritic cell activated** | **NA** | **NA** | **NA** | **NA** | ******* | **NA** | **NA** | **NA** |
| Neutrophil | NA | NA | ** | ns | ns | ns | NA | NA |
| NK cell | NA | NA | NA | ns | ns | ns | ns | NA |
| Plasmacytoid dendritic cell | NA | NA | NA | NA | ns | NA | NA | NA |
| stroma score | NA | ** | NA | NA | * | NA | NA | NA |
| T cell | NA | NA | NA | NA | NA | ns | NA | NA |
| T cell CD4+ | NA | NA | ns | ns | NA | NA | ns | NA |
| T cell CD4+ (non-regulatory) | NA | NA | NA | NA | ns | NA | NA | NA |
| T cell CD4+ central memory | NA | NA | NA | NA | ns | NA | NA | NA |
| T cell CD4+ effector memory | NA | NA | NA | NA | ns | NA | NA | NA |

| | | | | | | | | |
|---|---|---|---|---|---|---|---|---|
| T cell CD4+ memory | NA | NA | NA | NA | ns | NA | NA | NA |
| T cell CD4+ naive | NA | NA | NA | NA | ns | NA | NA | NA |
| T cell CD4+ Th1 | NA | NA | NA | NA | ns | NA | NA | NA |
| T cell CD4+ Th2 | NA | NA | NA | NA | ns | NA | NA | NA |
| T cell CD8+ | NA | NA | ns | ns | ns | ns | * | NA |
| T cell CD8+ central memory | NA | NA | NA | NA | ns | NA | NA | NA |
| T cell CD8+ effector memory | NA | NA | NA | NA | ns | NA | NA | NA |
| T cell CD8+ naive | NA | NA | NA | NA | ns | NA | NA | NA |
| T cell gamma delta | NA | NA | NA | ns | ns | NA | NA | NA |
| T cell NK | NA | NA | NA | NA | ns | NA | NA | NA |
| **T cell regulatory (Tregs)** | **NA** | **NA** | **NA** | **ns** | ******* | **NA** | **NA** | **NA** |
| **tumor purity fraction** | ******** | **NA** | **NA** | **NA** | **NA** | **NA** | **NA** | **NA** |
| **tumor purity score** | **NA** | ****** | **NA** | **NA** | **NA** | **NA** | **NA** | **NA** |
| uncharacterized cell | NA | NA | NA | NA | NA | NA | ns | NA |

**Table S5 – Bacterial counts impacting specific immune cells across the seven methods that score immune cell infiltration, excluding paired normal samples from the control group.**

NA = Not applicable because the method does not report that type of immune cell. ns = not significant. * = p<0.05, ** = p<0.01, *** p<0.001, **** p<0.0001. Statistical comparison was carried out with Ordinary Least Squares regression to determine which independent variables (sample type, bacterial count, read count) explain the dependent outcome variable (the immune score). Highlighted rows with bold text indicate immune cell sub-populations only affected in CRC tissue by bacterial counts. A p-value ≤ 0.05 was considered statistically significant.

| Cell_type | Estimate_frac | Estimate_score | Timer | consensus_tme | xcell | mcp_counter | epic | quantiseq |
|---|---|---|---|---|---|---|---|---|
| B cell | NA | NA | ns | ns | ns | ns | ns | NA |
| B cell memory | NA | NA | NA | NA | ns | NA | NA | NA |
| B cell naive | NA | NA | NA | NA | ns | NA | NA | NA |
| B cell plasma | NA | NA | NA | NA | ns | NA | NA | NA |
| Cancer associated fibroblast | NA | NA | NA | ns | ns | ns | ns | NA |
| Class-switched memory B cell | NA | NA | NA | NA | ns | NA | NA | NA |
| Common lymphoid progenitor | NA | NA | NA | NA | ns | NA | NA | NA |
| Common myeloid progenitor | NA | NA | NA | NA | ns | NA | NA | NA |
| cytotoxicity score | NA | NA | NA | ns | NA | ns | NA | NA |
| Endothelial cell | NA | NA | NA | ns | ns | ns | ns | NA |
| Eosinophil | NA | NA | NA | ns | ns | NA | NA | NA |
| Granulocyte-monocyte progenitor | NA | NA | NA | NA | ns | NA | NA | NA |
| Hematopoietic stem cell | NA | NA | NA | NA | ns | NA | NA | NA |
| immune score | NA | ns | NA | ns | ns | NA | NA | NA |
| Macrophage | NA | NA | ns | ns | ns | NA | ns | NA |
| Macrophage M1 | NA | NA | NA | ns | ns | NA | NA | NA |
| Macrophage M2 | NA | NA | NA | ns | ns | NA | NA | NA |
| Macrophage/Monocyte | NA | NA | NA | NA | NA | ns | NA | NA |
| Mast cell | NA | NA | NA | NA | ns | NA | NA | NA |
| MAST cell | NA | NA | NA | ns | NA | NA | NA | NA |
| microenvironment score | NA | NA | NA | NA | ns | NA | NA | NA |
| Monocyte | NA | NA | NA | ns | ns | ns | NA | NA |
| Myeloid dendritic cell | NA | NA | ns | ns | ns | ns | NA | NA |
| Myeloid dendritic cell activated | NA | NA | NA | NA | ns | NA | NA | NA |
| Neutrophil | NA | NA | ns | ns | ns | ns | NA | NA |
| NK cell | NA | NA | NA | ns | ns | ns | ns | NA |
| Plasmacytoid dendritic cell | NA | NA | NA | NA | ns | NA | NA | NA |
| stroma score | NA | ns | NA | NA | ns | NA | NA | NA |
| T cell | NA | NA | NA | NA | NA | ns | NA | NA |
| T cell CD4+ | NA | NA | ns | ns | NA | NA | ns | NA |
| T cell CD4+ (non-regulatory) | NA | NA | NA | NA | ns | NA | NA | NA |
| T cell CD4+ central memory | NA | NA | NA | NA | ns | NA | NA | NA |
| T cell CD4+ effector memory | NA | NA | NA | NA | ns | NA | NA | NA |
| T cell CD4+ memory | NA | NA | NA | NA | ns | NA | NA | NA |

| | | | | | | | | |
|---|---|---|---|---|---|---|---|---|
| T cell CD4+ naive | NA | NA | NA | NA | ns | NA | NA | NA |
| T cell CD4+ Th1 | NA | NA | NA | NA | ns | NA | NA | NA |
| T cell CD4+ Th2 | NA | NA | NA | NA | ns | NA | NA | NA |
| T cell CD8+ | NA | NA | ns | ns | ns | ns | ns | NA |
| T cell CD8+ central memory | NA | NA | NA | NA | ns | NA | NA | NA |
| T cell CD8+ effector memory | NA | NA | NA | NA | ns | NA | NA | NA |
| T cell CD8+ naive | NA | NA | NA | NA | ns | NA | NA | NA |
| T cell gamma delta | NA | NA | NA | ns | ns | NA | NA | NA |
| T cell NK | NA | NA | NA | NA | ns | NA | NA | NA |
| T cell regulatory (Tregs) | NA | NA | NA | ns | ns | NA | NA | NA |
| tumor purity fraction | ns | NA | NA | NA | NA | NA | NA | NA |
| tumor purity score | NA | ns | NA | NA | NA | NA | NA | NA |
| uncharacterized cell | NA | NA | NA | NA | NA | NA | ns | NA |

**Table S6 – *Bacteroides fragilis* impacting specific immune cells across the seven methods that score immune cell infiltration excluding paired normal samples from the control group.**

NA = Not applicable because the method does not report that type of immune cell. ns = not significant. * = $p<0.05$, ** = $p<0.01$, *** $p<0.001$. Statistical comparison was carried out with Ordinary Least Squares regression to determine which independent variables (sample type, bacterial count, read count) explain the dependent outcome variable (the immune score). Highlighted rows indicate immune cell sub-populations only affected in CRC tissue by *B. fragilis*. A p-value ≤ 0.05 was considered statistically significant.

| Cell_type | Estimate_frac | Estimate_score | Timer | consensus_tme | xcell | mcp_counter | epic | quantiseq |
|---|---|---|---|---|---|---|---|---|
| B cell | NA | NA | ns | ns | ns | ns | ns | NA |
| B cell memory | NA | NA | NA | NA | ns | NA | NA | NA |
| B cell naive | NA | NA | NA | NA | ns | NA | NA | NA |
| B cell plasma | NA | NA | NA | NA | ns | NA | NA | NA |
| Cancer associated fibroblast | NA | NA | NA | ns | ns | ns | ns | NA |
| Class-switched memory B cell | NA | NA | NA | NA | ns | NA | NA | NA |
| Common lymphoid progenitor | NA | NA | NA | NA | ns | NA | NA | NA |
| Common myeloid progenitor | NA | NA | NA | NA | ns | NA | NA | NA |
| cytotoxicity score | NA | NA | NA | ns | NA | ns | NA | NA |
| Endothelial cell | NA | NA | NA | ns | ns | ns | ns | NA |
| Eosinophil | NA | NA | NA | ns | ns | NA | NA | NA |
| Granulocyte-monocyte progenitor | NA | NA | NA | NA | ns | NA | NA | NA |
| Hematopoietic stem cell | NA | NA | NA | NA | ns | NA | NA | NA |
| immune score | NA | ns | NA | ns | ns | NA | NA | NA |
| Macrophage | NA | NA | ns | ns | ns | NA | ns | NA |
| Macrophage M1 | NA | NA | NA | ns | ns | NA | NA | NA |
| Macrophage M2 | NA | NA | NA | ns | ns | NA | NA | NA |
| Macrophage/Monocyte | NA | NA | NA | NA | NA | ns | NA | NA |
| Mast cell | NA | NA | NA | NA | ns | NA | NA | NA |
| MAST cell | NA | NA | NA | ns | NA | NA | NA | NA |
| microenvironment score | NA | NA | NA | NA | ns | NA | NA | NA |
| Monocyte | NA | NA | NA | ns | ns | ns | NA | NA |
| Myeloid dendritic cell | NA | NA | ns | ns | ns | ns | NA | NA |
| Myeloid dendritic cell activated | NA | NA | NA | NA | ns | NA | NA | NA |
| Neutrophil | NA | NA | ns | ns | ns | ns | NA | NA |
| NK cell | NA | NA | NA | ns | ns | ns | ns | NA |
| Plasmacytoid dendritic cell | NA | NA | NA | NA | ns | NA | NA | NA |
| stroma score | NA | ns | NA | NA | ns | NA | NA | NA |
| T cell | NA | NA | NA | NA | NA | ns | NA | NA |
| T cell CD4+ | NA | NA | ns | ns | NA | NA | ns | NA |
| T cell CD4+ (non-regulatory) | NA | NA | NA | NA | ns | NA | NA | NA |
| T cell CD4+ central memory | NA | NA | NA | NA | ns | NA | NA | NA |
| **T cell CD4+ effector memory** | **NA** | **NA** | **NA** | **NA** | ***** | **NA** | **NA** | **NA** |
| T cell CD4+ memory | NA | NA | NA | NA | ns | NA | NA | NA |

| | | | | | | | | |
|---|---|---|---|---|---|---|---|---|
| T cell CD4+ naive | NA | NA | NA | NA | ns | NA | NA | NA |
| T cell CD4+ Th1 | NA | NA | NA | NA | ns | NA | NA | NA |
| T cell CD4+ Th2 | NA | NA | NA | NA | ns | NA | NA | NA |
| T cell CD8+ | NA | NA | ns | ns | ns | ns | ns | NA |
| T cell CD8+ central memory | NA | NA | NA | NA | ns | NA | NA | NA |
| T cell CD8+ effector memory | NA | NA | NA | NA | ns | NA | NA | NA |
| T cell CD8+ naive | NA | NA | NA | NA | ns | NA | NA | NA |
| T cell gamma delta | NA | NA | NA | ns | ns | NA | NA | NA |
| T cell NK | NA | NA | NA | NA | ns | NA | NA | NA |
| T cell regulatory (Tregs) | NA | NA | NA | ns | ns | NA | NA | NA |
| tumor purity fraction | ns | NA | NA | NA | NA | NA | NA | NA |
| tumor purity score | NA | ns | NA | NA | NA | NA | NA | NA |
| uncharacterized cell | NA | NA | NA | NA | NA | NA | ns | NA |

**Table S7 – *Fusobacterium nucleatum* impacting specific immune cells across the seven methods that score immune cell infiltration excluding paired normal samples from the control group.**

NA = Not applicable because the method does not report that type of immune cell. ns = not significant. * = $p<0.05$. Statistical comparison was carried out with Ordinary Least Squares regression to determine which independent variables (sample type, bacterial count, read count) explain the dependent outcome variable (the immune score). Highlighted rows indicate immune cell sub-populations only affected in CRC tissue by *F. nucleatum*. A p-value ≤ 0.05 was considered statistically significant.

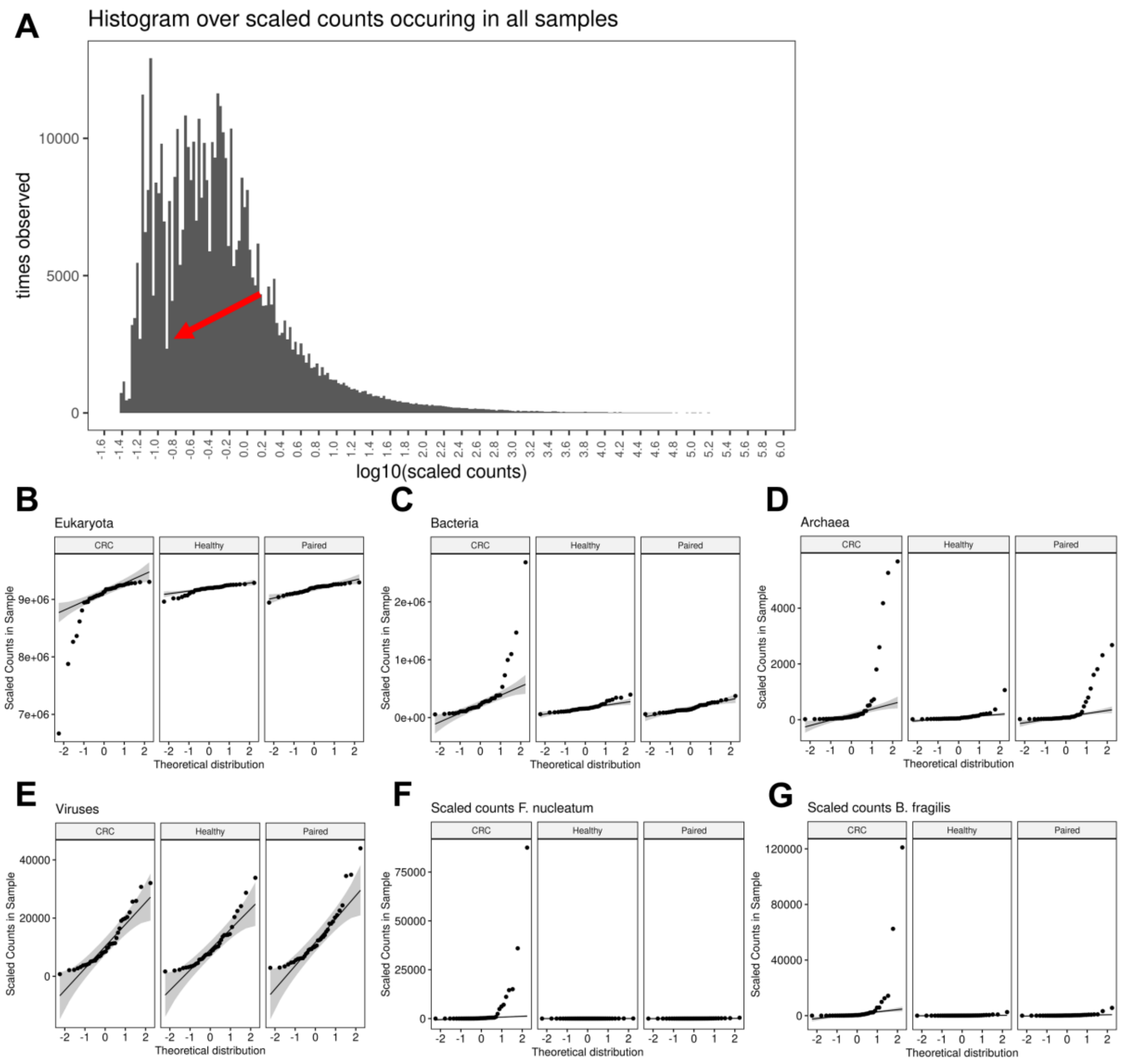


**Figure S1 - Noise filtering and distribution of counts assigned to Eukaryota, bacteria, archaea, viruses, *Fusobacterium nucleatum,* and *Bacteroides fragilis.***

**A**) Histogram showing the counts distribution over log-10 transformed scaled counts. The red arrow indicates the intersection between the populations, and all scaled counts < log-0.9 (indicated by red arrow) were set to 0 to remove noise. **B+C+D+E+F+G)** Normal distribution of scaled kingdom counts (y-axis) presented per sample across groups for Eukaryota (A), Bacteria (B), Archaea (C), virus (D), *F. nucleatum* (F), and *B. fragilis* (G).

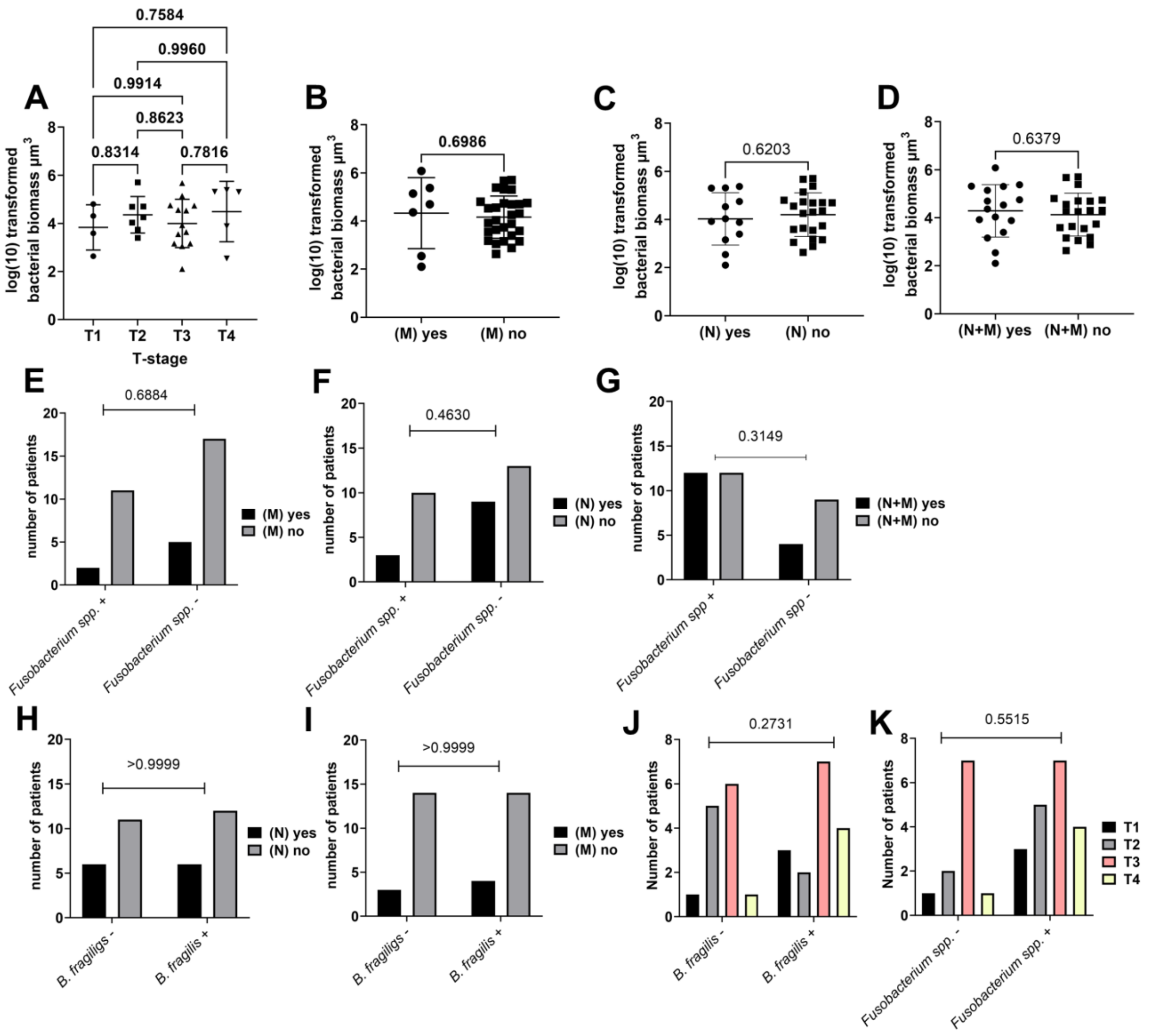


**Figure S2 - bacterial biomass was not associated with tumor staging, lymph node metastasis, or distant metastasis**

**A)** Logarithmic (log) transformed bacterial biomass measured in cubic micrometers ($\mu m^3$) according to tumor stage (T1-4). **B+C+D)** Log-transformed bacterial biomass measured in $\mu m^3$ according to distant (B) metastasis (M), Lymph node (C) metastasis (N), or both (D**).** All biomass measurements were measured with the Imaris software through thresholding of fluorescence intensity. **E+F+G)** Prevalence of *Fusobacterium spp.* compared with the number of patients with distant (E) metastasis (M), Lymph node (F) metastasis (N), or both (G**). H+I)** Prevalence of *Bacteroides fragilis* compared with the number of patients with lymph node (H) metastasis (N) or distant (I) metastasis (M**). J+K)** Number of patients and tumor staging (T1-4) compared to the prevalence of *B. fragilis* (J) and *Fusobacterium spp.* (K). Statistical comparison was carried out with one-way ANOVA **(A),** unpaired t-test **(B+C)**, Fisher's exact t-test **(A+B+C+D),** and chi-square test **(E+F)**. Bars represent standard deviation (SD); a p-value $\leq 0.05$ was considered statistically significant.

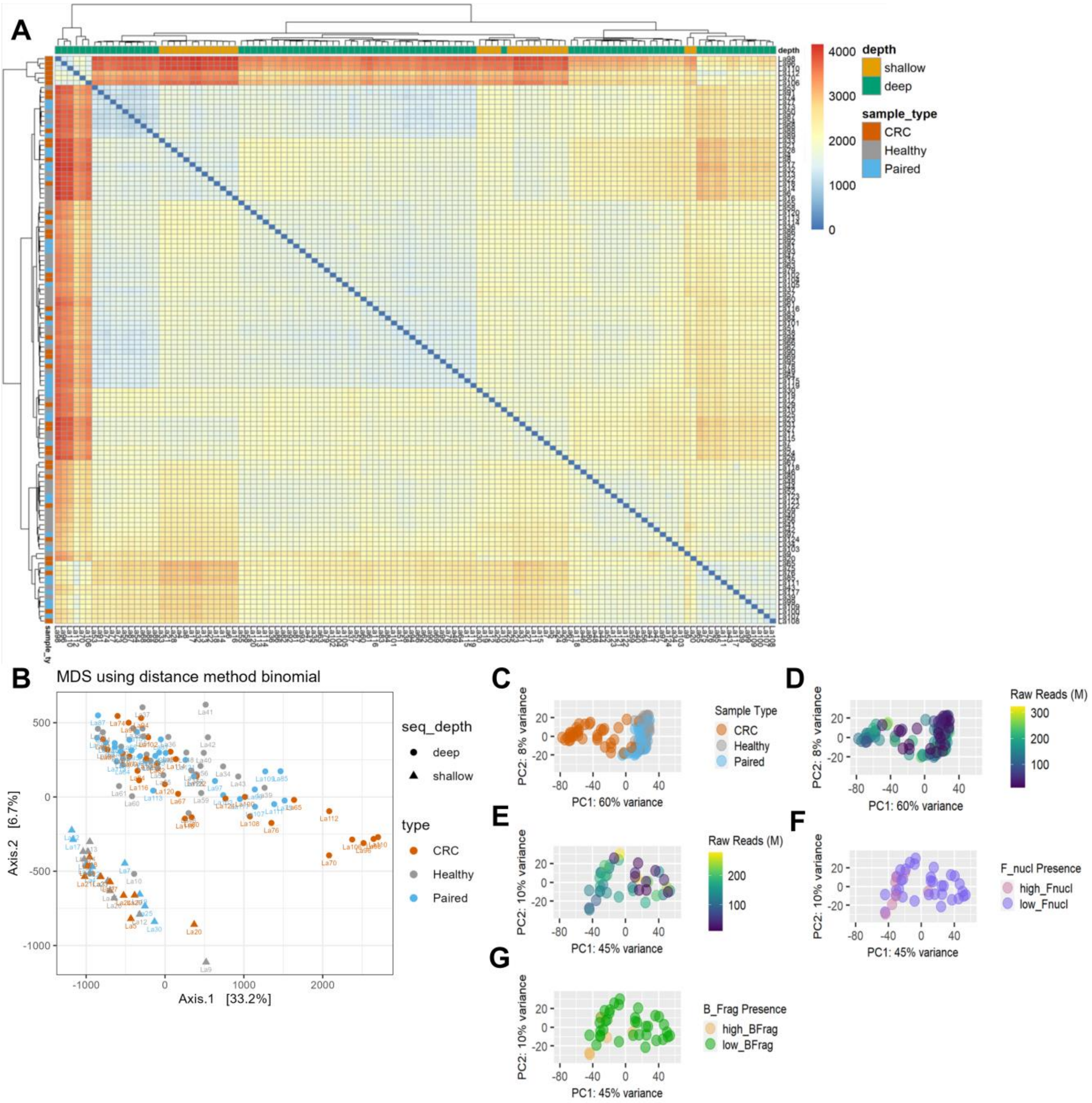


**Figure S3 - Principal-component analysis of normalized expression data.**

**A+B)** The clustering of CRC, healthy, and paired normal tissue samples according to sequencing depth (deep vs. shallow) is presented as a heatmap (A) and 2D scatterplot (B). **C)** All data is colored according to groups. **D)** All data is colored according to sequencing depth. **E)** CRC samples are colored by sequencing depth. **F)** CRC samples are colored according to the presence of *F. nucleatum*. Samples with high *F. nucleatum* counts were defined as those samples departing from the normal distribution in Figure S1, whereas low were those that followed the normal distribution. **G)** CRC samples are colored according to the presence of *B. fragilis*. Samples with high *B. fragilis* counts were defined as those departing from the normal distribution in Figure S1, whereas low were those following the normal distribution.

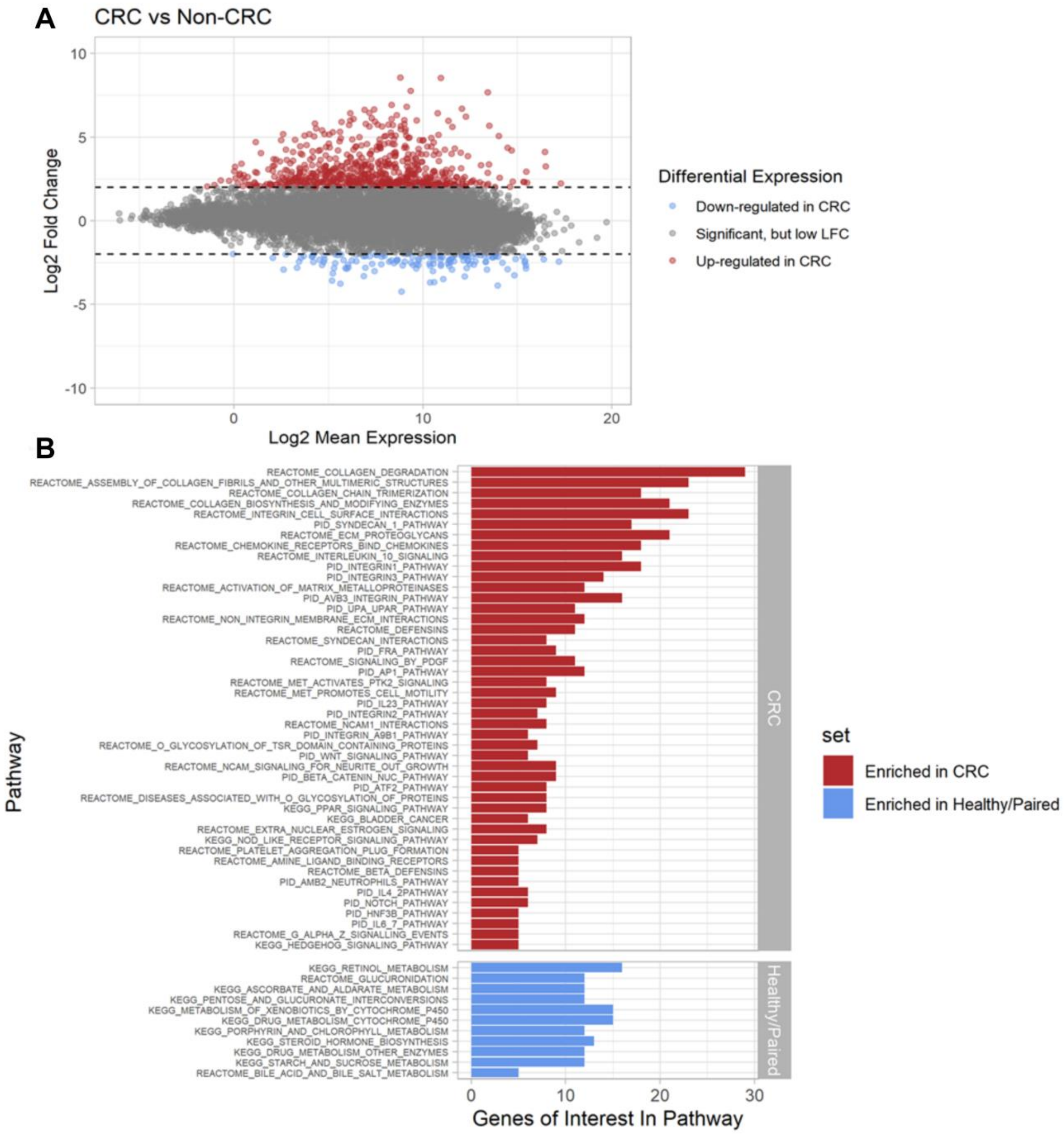


**Figure S4 - Differentially expressed genes and enriched biological pathways in CRC and non-CRC**

**A)** MA plot showing the distribution of significantly differentiated genes between CRC and Non-CRC (healthy and paired samples). Coloring highlights the 20 most significant DEGs with an adjusted p-value less than 0.05 and absolute log2 fold-change >2. Coloring highlights genes with an adjusted p-value less than 0.05 and absolute log2 fold-change >2. **B)** Pathways demonstrating significant enrichment of differentially expressed genes (Fishers exact test) for CRC or non-CRC. The Kyoto Encyclopedia of Genes and Genomes (KEGG), Pathway Interaction Database (PID), and REACTOME (a database of reactions, pathways, and biological processes) databases were used to identify pathways.

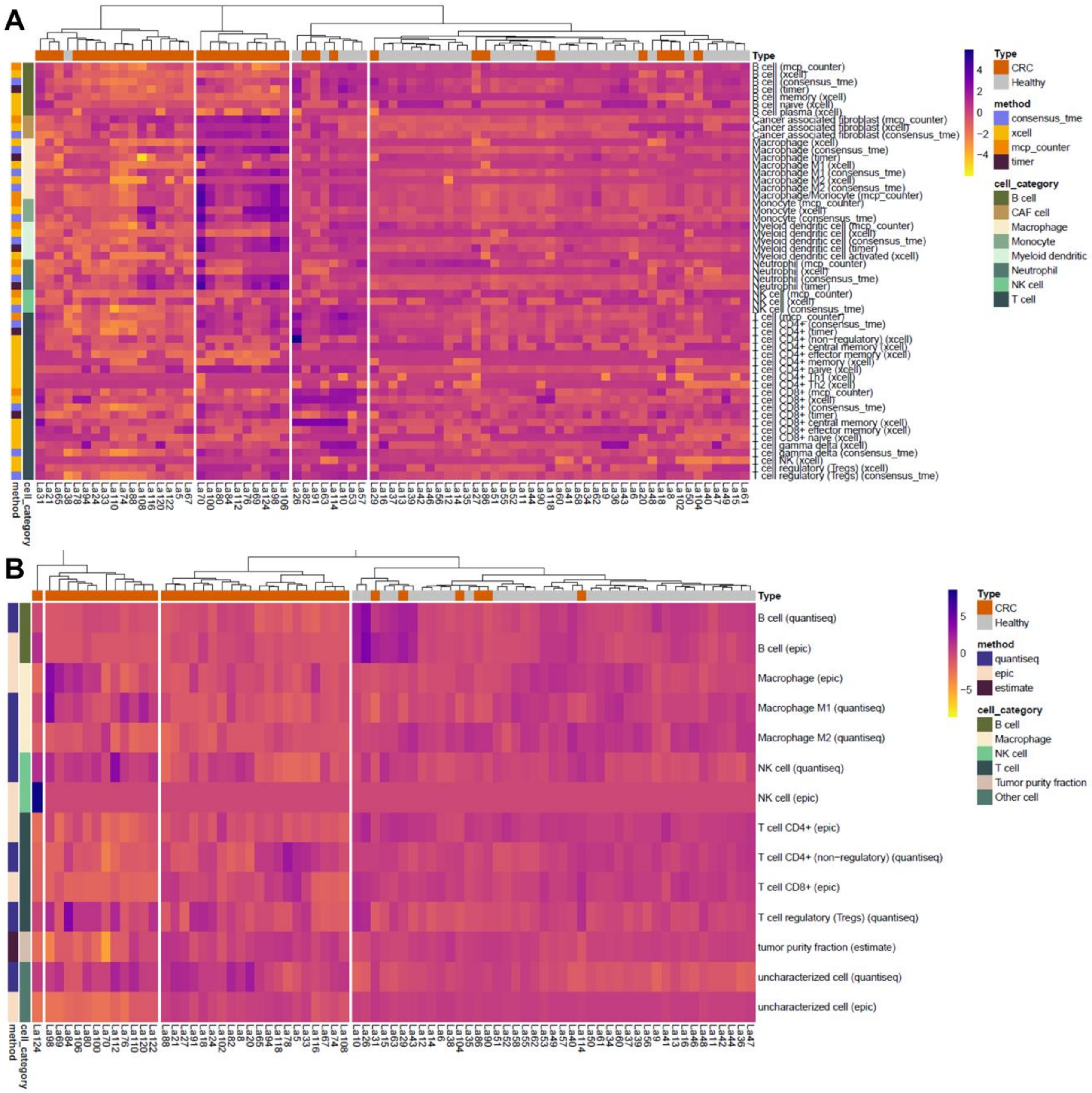


**Figure S5 - Clustering of samples according to immune cell infiltration 2**

**A+B)** Heatmaps showing immune cell profiles in CRC and healthy tissue, presented as normalized scores from the concensus_tme, xcell, mcp_counter, and timer immune scoring systems (A), and fractions from the quantisec, epic, and estimate immune scoring systems (B). Coloring from yellow (-4) to purple (4) indicates the degree of infiltration, where purple is high infiltration.
